# The neuropathy-causing *GARS1*^Δ*ETAQ*^ mutation drives pathology in subsets of motor and sensory neurons in mice

**DOI:** 10.64898/2026.07.05.736541

**Authors:** Rebecca L. Simkin, Aurélie Paulo-Ramos, Qiuhan Lang, Elena R. Rhymes, Sunaina Surana, David Villarroel-Campos, Sijiang Liu, Roberto Bellanti, Elena Veleva, Vlada Drotsevitch, Owen Swann, Amanda Heslegrave, Henrik Zetterberg, Michael P. Lunn, Robert W. Burgess, James N. Sleigh

**Author notes:** Equal contribution (Rebecca L. Simkin, Aurélie Paulo-Ramos).

## Abstract

Charcot-Marie-Tooth disease type 2D (CMT2D) results from gain-of-function mutations in *GARS1*, which encodes glycyl-tRNA synthetase (GlyRS), the enzyme responsible for charging transfer RNA (tRNA) with glycine. There are several CMT2D mouse models, but *Gars*^Δ*ETAQ*/+^ is the only one that bears a patient-sourced mutation. Created using CRISPR/Cas9 to model a 12-nucleotide *de novo GARS1* deletion identified in an unusually severe CMT2D patient, *Gars*^*Δ*ETAQ*/+*^ mice have previously been shown to display several neuromuscular phenotypes; motor axon loss, denervated neuromuscular junctions (NMJs) and reduced muscle function. Here, we extend these analyses to provide a more comprehensive understanding of both motor and sensory nerve deficits across hind- and fore-limbs. At 3 months, *Gars*^Δ*ETAQ*/+^ mice possess sex-independent alterations in the levels of neuropathy biomarkers – including decreased NfL and increased periaxin – alongside reduced muscle endurance and strength, and impairments in the sensory modalities of mechanosensation, proprioception and nociception. Underpinning these dysfunctions, we identified site-specific defects comprising altered sensory neuron populations, muscle spindle loss, reduced motor neuron size, disrupted NMJ innervation and maturation, and reduced axonal transport of signalling endosomes *in vivo*. Together, these experiments show that *Gars*^Δ*ETAQ*/+^ mice display robust and selective peripheral nerve pathology that manifests in a general distal-to-proximal fashion, priming this CMT2D allele for testing treatments and evaluating mechanisms underlying peripheral nerve vulnerability.

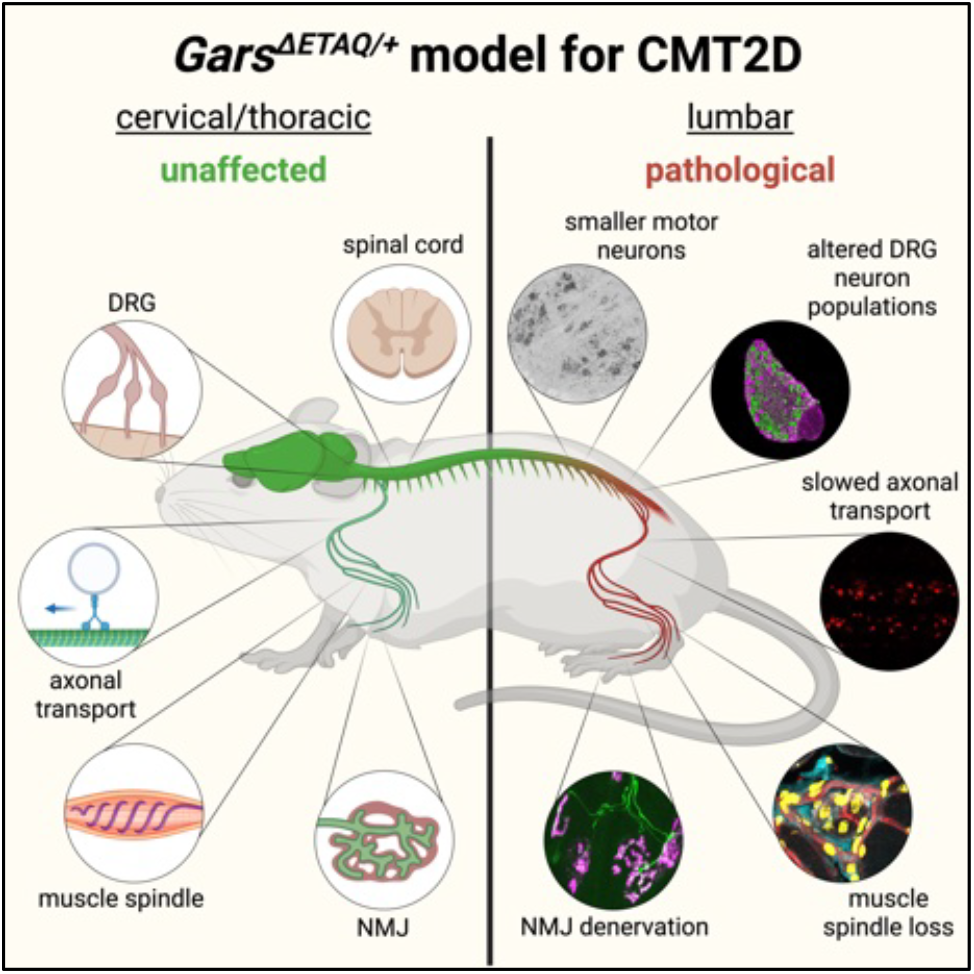

## Introduction

Charcot-Marie-Tooth disease (CMT) is the most common inherited neuromuscular condition (Deenen et al., 2025). Patients display motor and sensory nerve dysfunction that leads to muscle weakness and atrophy, musculoskeletal deformities, absent deep tendon reflexes and sensory deficits (Reilly et al., 2011, Burns et al., 2026). These symptoms usually manifest during adolescence, thus cause life-long disability, and predominantly affect peripheral nerves innervating distal parts of the body, *i*.*e*., primarily the feet and then the hands. CMT can be broadly divided into two main subtypes: type 1/demyelinating CMT, which results from primary Schwann cell dysfunction that reduces nerve conduction velocities (NCVs), and type 2/axonal CMT, characterised by axon degeneration without clear conduction impairment. Symptom support, surgical interventions and rehabilitative approaches are currently the only available treatment options, while the slowly progressive nature and genetic diversity of the disease complicate CMT clinical trials (Rossor et al., 2020). Nevertheless, several non-specific blood biomarkers for CMT, including neurofilament light chain (NfL) and growth differentiation factor 15 (GDF15), have been recently identified that may aid in this endeavour, at least for some neuropathy subtypes (Millere et al., 2021, Rossor et al., 2022, Jennings et al., 2022).

CMT and the closely related motor- and sensory-specific neuropathies result from mutations in more than 130 different genes, which encode proteins with a diversity of functions (Pipis et al., 2019, Record et al., 2024). The largest gene family linked to CMT are the aminoacyl-tRNA synthetases (ARSs), essential enzymes that charge amino acids to their cognate transfer RNAs (tRNAs) for protein synthesis (Wei et al., 2019, Burgess and Storkebaum, 2023). *GARS1* encodes glycyl-tRNA synthetase (GlyRS) and was the first ARS gene linked to neuropathy, with mutations causing both CMT subtype 2D (CMT2D; OMIM #601472) and distal hereditary motor neuropathy type V (dHMNV; OMIM #600794), the latter selectively affecting motor neurons (Antonellis et al., 2003). As a consequence, CMT2D is the best-studied ARS neuropathy and has several mouse models available to investigate the pathomechanisms linking dominantly inherited *GARS1* mutations to the selective perturbation of peripheral nerve function (Kalotay et al., 2023). The three principal mutant *Gars* strains used to model CMT2D are the spontaneous *Gars*^*Nmf249/+*^ mutant (most severe) (Seburn et al., 2006), the mutagen-induced *Gars*^*C201R/+*^ (least severe) (Achilli et al., 2009), and the CRISPR/Cas9-generated *Gars*^Δ*ETAQ*/+^ (intermediate) (Morelli et al., 2019). Modelling CMT mutations in mice does not always result in a clear peripheral nerve phenotype (Juneja et al., 2019, Bosco et al., 2021); nonetheless, the mutant *Gars* alleles all display robust distal motor impairments that recapitulate aspects of the human disease – for example, reduced muscle mass and strength without motor neuron loss. Consequently, these mice have been instrumental in determining that CMT2D results from a toxic gain-of-function in GlyRS (Motley et al., 2011), mediated through neomorphic interactions that impair protein synthesis, activate the integrated stress response, perturb axonal transport, and disrupt key neuronal signalling pathways (He et al., 2011, He et al., 2015, Benoy et al., 2018, Mo et al., 2018, Spaulding et al., 2021, Zuko et al., 2021, Sleigh et al., 2023).

The *Gars*^Δ*ETAQ*/+^ strain models a severe 12-nucleotide *de novo* deletion in exon 8 of *GARS1*, which removes four highly conserved amino acids in GlyRS (glutamic acid 299 to glutamine 302). This ETAQ deletion was identified in a heterozygous 13 month-old female, who presented with impaired motor function and regression of motor milestones in both her legs and arms (Morelli et al., 2019). *Gars*^Δ*ETAQ*/+^ mice were generated using CRISPR/Cas9 genome-editing technology and display a clear peripheral neuropathy phenotype. By 12 weeks, the CMT2D strain presents with reduced body weight, impaired muscle function, a decline in motor axon number in the femoral nerve, diminished sciatic NCVs and disrupted neuromuscular junction (NMJ) innervation in hind-limb plantaris muscles (Morelli et al., 2019). Follow-up studies have identified additional *Gars*^Δ*ETAQ*/+^ phenotypes including reduced compound muscle action potentials in the gastrocnemius (Spaulding et al., 2021, Zuko et al., 2021), perturbed *in vivo* transport of signalling endosomes in motor axons (Sleigh et al., 2023), reduced myelin thickness in the sciatic nerve and abnormal gastrocnemius muscle histochemistry (Ozes et al., 2021).

However, several questions regarding the utility of the *Gars*^Δ*ETAQ*/+^ strain remain. For example, are blood biomarkers identified in *Gars*^*Nmf249/+*^ and *Gars*^*C201R/+*^ also altered in this model? Is the sensory nervous system impacted? Do motor neurons show a similar pattern of selective vulnerability to that observed in other CMT2D alleles? We therefore set out to address these gaps in understanding, with a view to providing a more detailed picture of the *Gars*^Δ*ETAQ*/+^ phenotype, which will facilitate insights into underlying pathomechanisms. Moreover, by evaluating peripheral nerves innervating the hind- and fore-limbs, we aimed to determine whether the upper-limb predominance reported across most CMT2D/dHMNV patients is present in *Gars*^Δ*ETAQ*/+^ mutants (Sivakumar et al., 2005, McMacken et al., 2021).

## Materials and Methods

### 2.1 Ethical Approval

Experiments involving mice were performed under license from the UK Home Office in accordance with the Animals (Scientific Procedures) Act (1986) and were approved by the UCL Queen Square Institute of Neurology Ethical Review Committee.

### 2.2 Animals

Mice were maintained under a 12 h light/dark cycle at a constant room temperature of ≈21°C with ad libitum access to water and Teklad Global 18% Protein Rodent Diet (Inotiv, 2018C). As standard, cages were enriched with nesting material, plastic/cardboard tubes and wooden chew sticks. *Gars*^Δ*ETAQ*/+^ (MGI: 6393282) and *Gars*^*C201R/+*^ (MGI: 3760297) mice were maintained as male heterozygous mutants × female wild-type breeding pairs on the C57BL/6J background. Both female and male mice were assessed (see **Supplementary Table 1** for details). Multiple experiments were performed on individual mice whenever possible to reduce the overall number of animals used. Mice were sacrificed at approximately 3 months of age, unless otherwise stated (see **Supplementary Table 1**). Post-natal day 1 (P1) was defined as the day following the first observation of a litter. Genotyping from ear biopsies was performed by Transnetyx (Cordova, TN, USA) using custom-made probes.

### 2.3 Body Weight

Mice were weighed using standard laboratory scales. Relative body weight was calculated by expressing each raw value as a percentage of the sex-matched wild-type average.

### 2.4 Blood Sampling and Analysis

Mice were euthanised using an overdose of pentobarbital sodium, and blood was collected via transcardiac puncture. Sampled blood was separated into both EDTA-lined tubes and 1.5 ml microcentrifuge tubes. Blood in EDTA-lined tubes was immediately mixed by swirling before transfer to a 1.5 ml microcentrifuge tube for centrifugation at 1,200 × g for 15 min at 4°C. Plasma was then collected, aliquotted and stored at -70°C. Blood in 1.5 ml microcentrifuge tubes was left at room temperature for 30-60 min before centrifugation at 3,500 × g for 15 min at 4°C. Serum was then collected, aliquotted and stored at -70°C. Single molecule array (Simoa) on an HD-X analyser (Quanterix) was performed to measure concentrations of NfL and periaxin in plasma, and peripherin in serum, as previously described (Rohrer et al., 2016, Keddie et al., 2023, Bellanti et al., 2025). Plasma Gdf15 was measured using a Mouse/Rat GDF-15 Quantikine ELISA Kit (R&D Systems, MGD150), following the manufacturer’s instructions.

### 2.5 Sensory Behaviour

Sensory function was assessed largely as previously described in *Gars*^*C201R/+*^ mice (Sleigh et al., 2017). Prior to sensory behavioural testing (see below), mice were transferred to the testing room and acclimatised for 45 min in a plastic chamber (10 × 10 × 14 cm) placed on an elevated wire grid.

#### 2.5.1 Von Frey Test

The glabrous skin of the hind-paw was stimulated using calibrated von Frey filaments (0.07 to 1.14 g) (Ugo Basile, 37450-275). Von Frey responses were first determined using the ‘up and down’ method (Bonin et al., 2014), starting at 0.16 g, to calculate the paw withdrawal score. The response frequency was then obtained by stimulating the hind-paw five times with each filament (0.07 to 1.14 g in ascending order) to determine the percentage paw withdrawal response. The stimulations were performed alternating between the left and right hind-paws, with at least 1 min between each stimulation.

#### 2.5.2 Balance Beam Test

Mice were trained for two consecutive days to cross balance beams of different widths (24, 12 and 6 mm). On the test day, mice were recorded crossing the 12 mm beam twice using a camera phone. The number of paw slips were then analysed using the recorded videos, as described previously (Carter et al., 2001).

#### 2.5.3 Pinprick Test

The glabrous skin of the hind-paws was stimulated using a 30G needle with gentle pressure and without skin penetration. This stimulation was repeated five times, with intervals of at least 1 min, alternating between the left and right paws. Pinprick responses were calculated as a percentage of paw withdrawal.

#### 2.5.4 Hargreaves Test

Mice were placed in a plastic chamber on a glass floor and heat was applied to the hind-paw, as first described by Hargreaves and colleagues (Hargreaves et al., 1988). The response to radiant heat was measured using an infrared heat stimulus from the plantar test apparatus (Ugo Basile, 37570), which measures paw withdrawal latency. The infrared intensity was adjusted to 30% so that control mice displayed a latency of 6-10 s. A cut-off time of 15 s was applied to avoid tissue damage.

### 2.6 Motor Function

Wire-hang tests were performed by placing mice upside down on an inverted grid and measuring latency to fall, with a maximum cut-off time of 2 min, as previously described (Spaulding et al., 2016). The maximum time achieved over three trials was recorded. If a mouse reached the 2 min cut-off in the first or second trial, no additional tests were performed. All-limb grip strength was assessed using a Grip Strength Meter (Bioseb, GS3), as previously described (Rhymes et al., 2024).

### 2.7 Immunofluorescence

Dorsal root ganglia (DRG) from spinal levels lumbar 1 (L1) to L5 and cervical 4 (C4) to C8 were dissected (Sleigh et al., 2016, Sleigh et al., 2020e), sectioned at 10 µm, immunostained and analysed as described elsewhere (Sleigh et al., 2017, Sleigh et al., 2020a). An average of 2,240 ± 100 lumbar cells (range: 1,780-2,712) and 1,717 ± 78 cervical cells (range: 1,298-2,407) across three DRG were assessed in each mouse. The percentages of neurofilament 200-positive (NF200+) and peripherin-positive (peripherin+) cells per mouse were calculated by averaging values from individual DRG. Cells with increased GlyRS expression (above average wild-type levels) were first identified visually in the single fluorescence channel and marked on the image using the Cell Counter tool in ImageJ (http://rsb.info.nih.gov/ij/). NF200-positive cells were then independently identified in the second fluorescence channel. These datasets were used to calculate the percentage of GlyRS-elevated cells that were positive for NF200, and the proportion of NF200+ cells lacking increased GlyRS expression. DRG sections used for GlyRS analysis were stained in parallel and imaged using the same confocal settings. Soleus and biceps muscles were dissected, sectioned at 20 µm, immunostained and analysed as detailed previously (Sleigh et al., 2017). Spinal cords were dissected from 4% paraformaldehyde-perfused mice and embedded in Tissue-Tek O.C.T. (Sakura Finetek, 4583). Lumbar and thoracic segments were then sectioned at 30 µm. Choline acetyltransferase (ChAT)-positive neurons within lumbar 3 (L3) to L5 and thoracic 4 (T4) to T6 anterior horns were counted and cell body areas measured as previously described (Sleigh et al., 2023, Rhymes et al., 2024). Epitrochleoanconeus (ETA), flexor digitorum brevis (FDB) and both hind- and fore-limb lumbrical muscles were dissected, processed for NMJ imaging, and analysed as previously described (Sleigh et al., 2014a, Mech et al., 2020, Villarroel-Campos et al., 2022, Simkin et al., 2025). One hundred NMJs per mouse were assessed to determine degenerative and maturation phenotypes. Primary and secondary antibodies used for immunofluorescence are listed in **Supplementary Table 2** and **3**, respectively. In addition, AlexaFluor555-conjugated α-bungarotoxin (α-BTX; Life Technologies, B35451) was used at 1:1,000 to identify post-synaptic acetylcholine receptors (AChRs). Fluorescent imaging of fixed samples and live nerves (see below) was performed on either an inverted LSM780 or LSM980 laser-scanning microscope (ZEISS).

### 2.8 Protein Extraction and Western Blotting

DRG and peripheral nerves (sciatic, median and ulnar) were dissected from PBS-perfused and non-perfused mice. Proteins were extracted from tissues and evaluated by western blotting as detailed previously (Sleigh et al., 2023). Primary and secondary antibodies used for western blotting are listed in **Supplementary Table 2** and **3**, respectively. Densitometry was performed using total protein stained with 0.1% Coomassie Brilliant Blue R-250 (Thermo Fisher, 20278) as the loading control. Hook1 bands at ≈110 kDa and ≈90 kDa were quantified together in nerve analyses.

### 2.9 In Vivo Imaging of Signalling Endosome Axonal Transport

Live imaging of signalling endosome axonal transport was performed using an atoxic binding fragment of the tetanus neurotoxin (H_C_T; residues 875-1,315 fused to a cysteine-rich tag and a human influenza haemagglutinin epitope) labelled with Alexa Fluor 555 C2 Maleimide (Thermo Fisher Scientific, A20346), as formerly described (Gibbs et al., 2016, Sleigh et al., 2020d, Tosolini et al., 2021). Briefly, to assess transport in the hind-limbs, 3-5 µg H_C_T-555 was injected under isoflurane-induced anaesthesia into the tibialis anterior on one side of the body and the contralateral gastrocnemius; this enabled transport assessment in peripheral nerves supplying distinct hind-limb muscles of the same mouse. The side of injection (tibialis anterior versus gastrocnemius) was alternated between mice to eliminate time-under-anaesthesia and right-left biases. Imaging was performed 4-8 h later in both sciatic nerves. To assess transport in the fore-limbs, 3-5 µg H_C_T was injected under anaesthesia into the left fore-paw before imaging the median and ulnar nerves 4-8 h later (Lang et al., 2023). Imaging was performed in a pre-warmed environmental chamber set to 38°C and images were acquired every ≈3 seconds using a 63× Plan-Apochromat oil immersion objective (Zeiss). By selecting thicker H_C_T-positive axons, evaluation of axonal transport was performed in motor axons (Sleigh et al., 2020c). Nodes of Ranvier were avoided due to their slowing effect on cargo trafficking (Tosolini et al., 2024). 15 endosomes per axon and at least three axons per mouse were manually tracked using the TrackMate plugin on ImageJ (Tinevez et al., 2017). An endosome was determined to have paused if it remained stationary (within < 0.3 µm) for two consecutive frames. Endosome frame-to-frame speeds are presented in frequency histograms. Endosome reversals were infrequent (<1% of all frame-to-frame movements) and recorded as positive values in the histograms.

### 2.10 Statistics

Data were assumed to be normally distributed unless evidence to the contrary was provided by the Kolmogorov-Smirnov test for normality, while equal variance between groups was assumed. The Bonferroni correction was applied to the Kolmogorov-Smirnov tests within each experiment. Normally distributed data were analysed using unpaired *t*-tests, one-way analysis of variance (ANOVA) tests or two-way ANOVA tests followed by Šídák’s multiple comparisons test or uncorrected Fisher’s least significant difference test. Non-normally distributed data were analysed using a Mann-Whitney *U* test or Kruskal-Wallis test followed by Dunn’s multiple comparisons test. Sample sizes, which were pre-determined using power calculations and previous experience (Sleigh et al., 2014b, Sleigh et al., 2017, Sleigh et al., 2020a, Sleigh et al., 2020b, Sleigh et al., 2023), are reported in figure legends and **Supplementary Table 1**, and represent biological replicates (*i.e*., individual animals). Means ± standard error of the mean are plotted for all graphs. All tests were two-sided and an α-level of *P* < 0.05 was used to determine significance. GraphPad Prism 10 software (version 10.6.0) was used for statistical analyses and figure production.

## Results

### 3.1 Gars^ΔETAQ/+^ mice display reduced weight and altered blood biomarkers

To provide a more detailed understanding of peripheral neuropathy in *Gars*^Δ*ETAQ*/+^ mice, we performed extensive phenotyping at 3 months of age. This timepoint was selected because pronounced neuropathology is evident by this stage in the well-characterised *Gars*^*Nmf249/+*^ and *Gars*^*C201R/+*^ CMT2D models (Seburn et al., 2006, Achilli et al., 2009, Sleigh et al., 2014b). We first confirmed a clear reduction in body weight for both female and male mutants (**Fig. 1A**), consistent with prior observations (Morelli et al., 2019, Spaulding et al., 2021, Tadenev et al., 2024, Rice et al., 2025). By comparing mutant weights relative to wild-type, we observed that *Gars*^Δ*ETAQ*/+^ males showed a greater percentage deficit than mutant females (**Fig. 1B**), hinting that males could be more severely affected.

**Figure 1.**
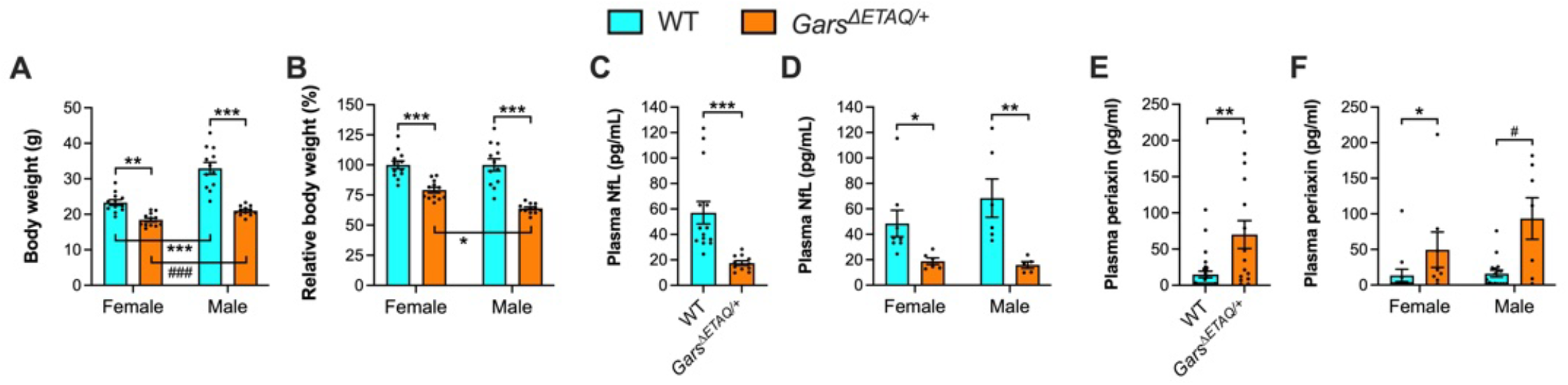
*Gars*^Δ*ETAQ*/+^ mice display reduced body weights and alterations in plasma NfL and periaxin levels. (**A-B**) Female and male *Gars*^Δ*ETAQ*/+^ mice display reduced body weight (A, genotype *P* < 0.001, sex *P* < 0.001, interaction *P* < 0.001 two-way ANOVA) and relative body weight (B, genotype *P* < 0.001, sex *P* = 0.026, interaction *P* = 0.026 two-way ANOVA) compared to wild-type. *n* = 12. (**C-D**) Neurofilament light chain (NfL) levels are diminished in *Gars*^Δ*ETAQ*/+^ mouse plasma (C, \*\*\**P* < 0.001 Mann-Whitney *U* test) in both females and males (D, genotype *P* < 0.001, sex *P* = 0.411, interaction *P* = 0.271 two-way ANOVA). *n* = 11-14 (C) and 5-8 (D). (**E-F**) In contrast, periaxin levels are increased in *Gars*^Δ*ETAQ*/+^ mouse plasma (E, \*\**P* = 0.002 Mann-Whitney *U* test) in both females and males (E, *P* = 0.009 Kruskal-Walli test). *n* = 15-29 (E) and 7-17 (F). For all graphs, data were generated from 3 month-old mice; \**P* < 0.05, \*\**P* < 0.01, \*\*\**P* < 0.001 Šídák’s/Dunn’s/Fisher’s post-hoc test (unless stated otherwise); ^#^*P* < 0.05 ^###^*P* < 0.001 unpaired *t*-test.

To provide an unbiased, systemic assessment of neuropathology, we evaluated levels of several blood biomarkers reported to be altered in CMT patients and/or mouse models: NfL (Millere et al., 2021, Rossor et al., 2022), periaxin (Bellanti et al., 2025), GDF15 (Jennings et al., 2022) and peripherin (Keddie et al., 2023). Similar to *Gars*^*C201R/+*^ mice (Rossor et al., 2022), NfL was decreased in plasma of *Gars*^Δ*ETAQ*/+^ mutants (**Fig. 1C**), and equally so in females and males (**Fig. 1D**). In contrast, periaxin levels were increased in both sexes (**Fig. 1E-F**), whereas *Gars*^*C201R/+*^ mice showed no change (**Supplementary Fig. 1A**), supporting the notion that *Gars*^Δ*ETAQ*/+^ is a more severe allele. However, the previously identified elevation of Gdf15 in serum of 5 month-old *Gars*^*Nmf249/+*^ and *Gars*^*C201R/+*^ mice was not replicated in *Gars*^Δ*ETAQ*/+^ plasma at 3 months (**Supplementary Fig. 1B-C**). We were also unable to detect changes in serum peripherin levels from either *Gars*^Δ*ETAQ*/+^ or *Gars*^*C201R/+*^ mutants (**Supplementary Fig. S1D-E**). Together, these data reveal NfL and periaxin as useful blood biomarkers for objective evaluation of the *Gars*^Δ*ETAQ*/+^ allele.

### 3.2 Sensory and motor function are impaired in Gars^ΔETAQ/+^ mice

We have previously shown that *Gars*^*C201R/+*^ mice display sensory defects across various different modalities – mechanosensation and proprioception are diminished, whereas mechanical and thermal nociception are enhanced (Sleigh et al., 2017). We therefore performed a similar battery of sensory behavioural tests on combined female and male *Gars*^Δ*ETAQ*/+^ mice. Using a series of von Frey filaments, we observed a reduced frequency of response to stronger stimuli, indicative of impaired mechanosensation (**Fig. 2A**). In the beam-walking test, we saw a greater number of paw slips by *Gars*^Δ*ETAQ*/+^ mice as they crossed a 12 mm beam, consistent with a proprioception defect (**Fig. 2B**). Contrasting with *Gars*^*C201R/+*^ mice, we found a reduced paw withdrawal frequency in response to a noxious pinprick stimulus, signifying impaired mechanical nociception (**Fig. 2C**). Finally, we identified reduced latency to paw withdrawal in response to a noxious heat stimulus, supporting enhanced thermal nociception (**Fig. 2D**). Together, these data indicate that deficits in three of the four sensory modalities tested in *Gars*^*C201R/+*^ mice are replicated in *Gars*^Δ*ETAQ*/+^, with mechanical nociception being enhanced in the former, but impaired in the latter.

**Figure 2.**
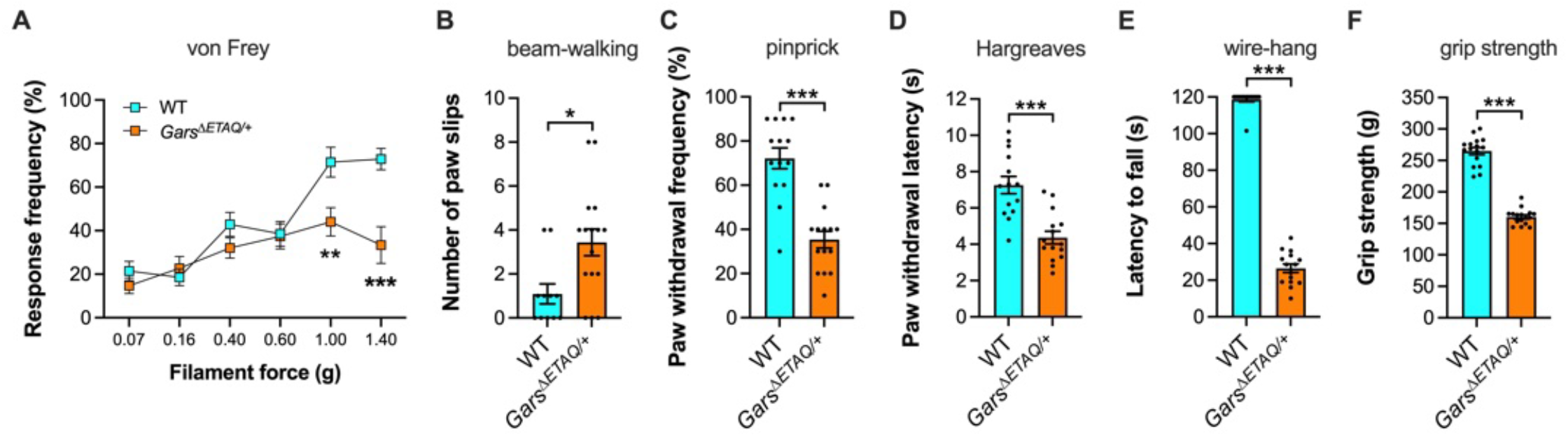
Sensory and motor function are impaired in *Gars*^Δ*ETAQ*/+^ mice. (**A-D**) *Gars*^Δ*ETAQ*/+^ mice display impairments in mechanosensation (A, genotype *P* < 0.001, force *P* < 0.001, interaction *P* = 0.001 two-way ANOVA; \*\**P* < 0.01, \*\*\**P* < 0.001 Šídák’s post-hoc test), proprioception (B, \**P* = 0.011 Mann-Whitney *U* test), mechanical nociception (C, \*\*\**P* < 0.001 unpaired *t*-test) and thermal nociception (D, \*\*\**P* < 0.001 unpaired *t*-test). (**E-F**) *Gars*^Δ*ETAQ*/+^ mice also display defective muscle endurance (E, \*\*\**P* < 0.001 Mann-Whitney *U* test) and maximal grip strength (F, \*\*\**P* < 0.001 unpaired *t*-test). For all graphs, data were generated from 3 month-old mice; *n* = 11-16.

Motor function deficits are well described across CMT2D models (Seburn et al., 2006, Achilli et al., 2009, Sleigh et al., 2017, Spaulding et al., 2021), including reduced latency to fall from an inverted wire grid and decreased grip strength in *Gars*^Δ*ETAQ*/+^ mice (Morelli et al., 2019, Zuko et al., 2021, Tadenev et al., 2024, Rice et al., 2025). To confirm that our colony has the expected motor phenotype, we replicated these prior observations by showing clear deficits in wire-hang (**Fig. 2E**) and grip strength (**Fig. 2F**), which confirm impaired muscle endurance and maximal strength, respectively.

We separated sensory and motor behavioural data by sex and observed that all deficits were similarly present in both female and male mutants, albeit with beam-walking impairments not reaching significance for either sex (**Supplementary Fig. 2**). This indicates that, despite differences in relative body weight, peripheral nerve function is not more severely affected in males. In summary, these data confirm that *Gars*^Δ*ETAQ*/+^ mice present with strong, sex-independent sensory and motor nerve dysfunction.

### 3.3 Gars^ΔETAQ/+^ lumbar but not cervical DRG display altered sensory neuron populations

Consistent with the sensory behavioural deficits, *Gars*^*C201R/+*^ mice display altered proportions of sensory neuron subtypes in the DRG, with fewer large-area, neurofilament 200-positive (NF200+) neurons (mainly mechanosensitive and proprioceptive), and an increase in smaller, peripherin+ neurons (primarily nociceptive) (Sleigh et al., 2017). This phenotype appears to be developmental in origin because it is present at birth (Sleigh et al., 2017), and is also selective, occurring in lumbar but not cervical DRG (Sleigh et al., 2020a). To determine whether *Gars*^Δ*ETAQ*/+^ mice display a similar defect, we dissected DRG from lumbar 1 (L1) to L5 and cervical 4 (C4) to C8 regions of the spinal column, and stained them with antibodies against NF200 and peripherin (**Fig. 3A**). Consistent with the *Gars*^*C201R/+*^ mutants, we observed a reduction in the percentage of NF200+ cells and an increase in peripherin+ cells in lumbar DRG (**Fig. 3B**), but not cervical (**Fig. 3C**). These findings complement western blotting data from lumbar DRG taken from 3-5 month-old *Gars*^Δ*ETAQ*/+^ mice (Sleigh et al., 2023). Furthermore, they are supported by western blot analysis on cervical DRG, showing no clear changes in the levels of NF200, peripherin or tropomyosin receptor kinase B (TrkB) receptor (**Supplementary Fig. 3A-B**). Binding to brain-derived neurotrophic factor (BDNF) and neurotrophin-4 (NT-4), TrkB is selectively found in a mechanosensitive sub-population of NF200+ neurons (Montano et al., 2010), and levels of the full-length receptor were previously found to be decreased in *Gars*^Δ*ETAQ*/+^ lumbar DRG (Sleigh et al., 2023).

**Figure 3.**
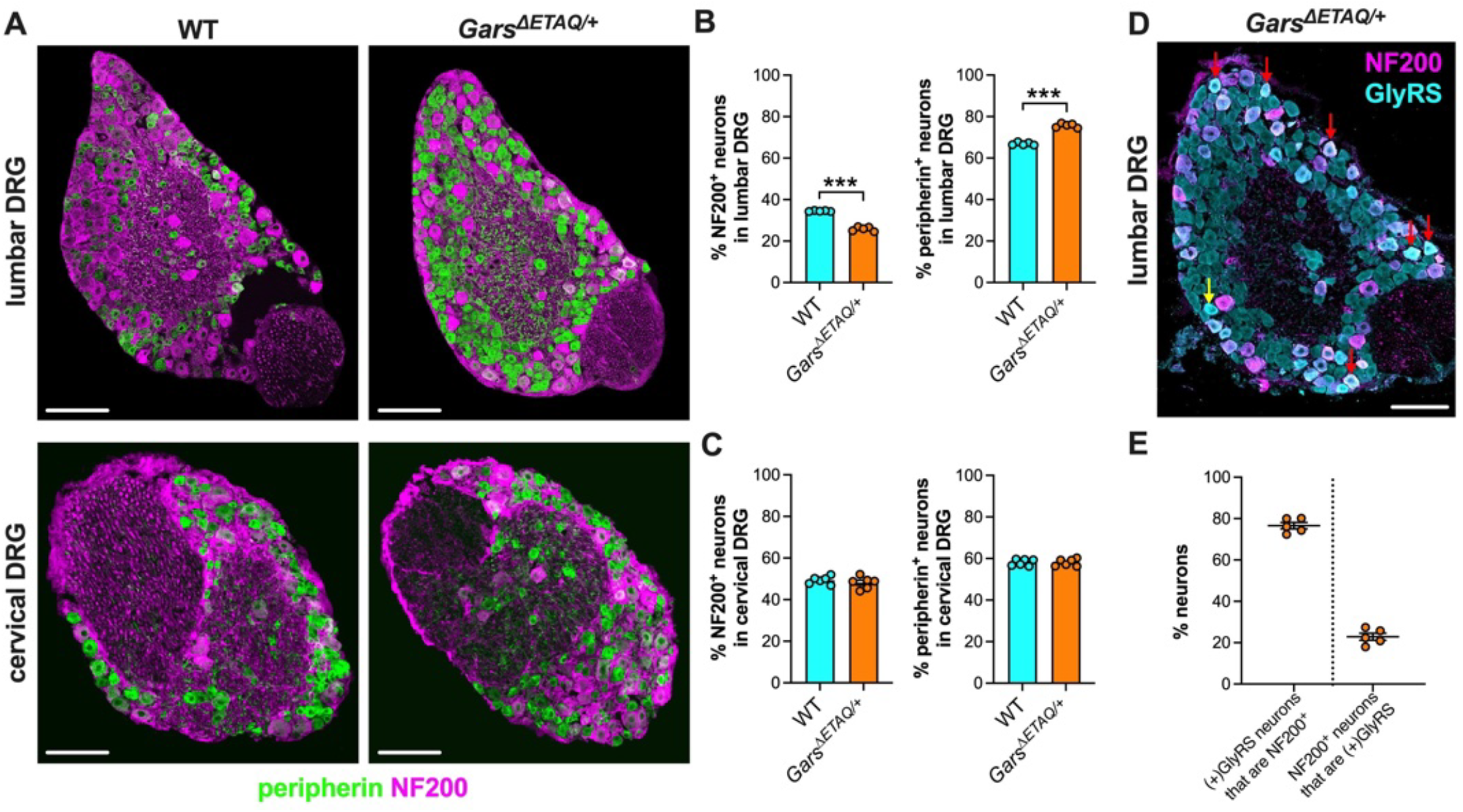
Sensory neuron subtypes are perturbed in *Gars*^Δ*ETAQ*/+^ lumbar DRG, but not cervical DRG. (**A**) Representative single-plane confocal images of wild-type (left) and *Gars*^Δ*ETAQ*/+^ (right) lumbar (top) and cervical (bottom) DRG sections stained for NF200 (magenta) and peripherin (green). (**B**) *Gars*^Δ*ETAQ*/+^ mice display a lower percentage of NF200^+^ neurons (left) and a higher percentage of peripherin^+^ neurons (right) in lumbar 1 (L1) to L5 DRG compared to wild-type. (**C**) The percentage of NF200^+^ neurons (left graph, *P* = 0.418 unpaired *t*-test) and peripherin^+^ neurons (right graph, *P* = 0.978 unpaired *t*-test) are unaffected in cervical 4 (C4) to C8 DRG of *Gars*^Δ*ETAQ*/+^ mice. (**D**) Representative single-plain confocal image of a *Gars*^Δ*ETAQ*/+^ lumbar DRG section stained for NF200 (magenta) and GlyRS (cyan). Red and yellow arrows highlight neurons with increased GlyRS that are NF200^+^ and NF200^-^, respectively. (**E**) The majority of neurons with increased GlyRS (designated as ‘(+)GlyRS’), express NF200; however, not all NF200^+^ cells exhibit increased GlyRS. For all panels, data were generated from 3 month-old mice; \*\*\**P* < 0.001 unpaired *t*-test; scale bars = 100 µm; *n* = 5-6.

In addition to altered sensory populations, we have previously found elevated GlyRS levels specifically within NF200+ neurons of lumbar DRG of *Gars*^*C201R/+*^ mice, the cause of which remains unknown (Sleigh et al., 2020a). Staining lumbar and cervical DRG from *Gars*^Δ*ETAQ*/+^ mice for both GlyRS and NF200 (**Fig. 3D, Supplementary Fig. 3C-D**), we replicated the selective increase in GlyRS (above normal wild-type levels) within NF200+ neurons in lumbar DRG (**Fig. 3E**), and further confirmed the absence of change in cervical DRG via immunofluorescence and western blot (**Supplementary Fig. 3A-B**). Together, these experiments confirm that *Gars*^Δ*ETAQ*/+^ mice display a perturbation in sensory neuron identity, as well as *Gars* expression, specifically within sensory neurons innervating hind-limbs.

### 3.4 Spindle pathology is present in Gars^ΔETAQ/+^ soleus but not biceps muscles

We have previously shown that *Gars*^*C201R/+*^ mice possess fewer muscle spindles in the soleus, and that the remaining proprioceptive nerve endings are poorly innervated (Sleigh et al., 2017). To determine whether this phenotype is also present in *Gars*^Δ*ETAQ*/+^ mutants and reflects the spatial selectivity of the DRG phenotype, we dissected hind-limb soleus and fore-limb biceps muscles, and stained sections with antibodies against synaptic vesicle protein 2 (SV2) and neurofilament (2H3) to identify spindles, and laminin to mark the basement membrane (**Fig. 4A-B**). We identified a clear reduction in spindle number and innervation in the soleus muscle (**Fig. 4C**), but not in the biceps (**Fig. 4D**), indicating a selective defect in proprioceptive nerve endings and supporting that hind-limbs are more affected than fore-limbs in *Gars*^Δ*ETAQ*/+^ mice.

**Figure 4.**
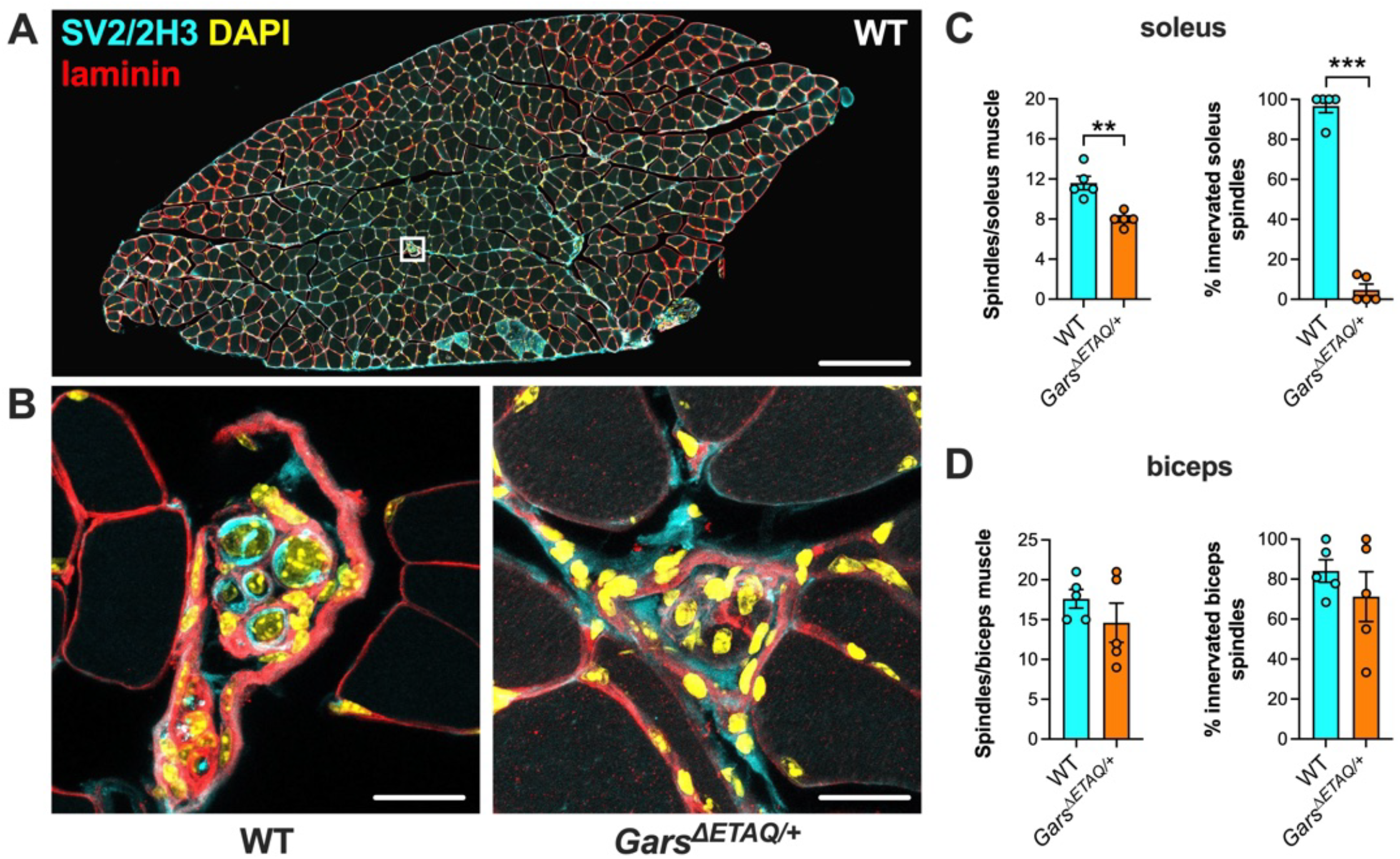
*Gars*^Δ*ETAQ*/+^ soleus muscles, but not biceps, have fewer and more severely denervated spindles. (**A**) Representative collapsed z-stack image of a wild-type soleus muscle section stained for SV2/2H3^+^ (cyan), laminin (red) and DAPI (yellow). The white square highlights a muscle spindle. Representative collapsed z-stack images of SV2/2H3^+^ muscle spindles from wild-type (left) and *Gars*^Δ*ETAQ*/+^ (right) soleus muscle sections. (**C**) *Gars*^Δ*ETAQ*/+^ mice have fewer spindles per soleus muscle (left, \*\**P* < 0.01), and mutant spindles display almost complete denervation (right, *P* = 0.008 Mann-Whitney *U* test). (**D**) Spindle number (left, *P* = 0.303) and innervation (right, *P* = 0.375) are unaffected in *Gars*^Δ*ETAQ*/+^ biceps muscles. For all panels, data were generated from 3 month-old mice and analysed by unpaired *t*-test (unless stated otherwise); scale bars = 200 (A) and 20 (B) µm; *n* = 5.

### 3.5 Motor neuron size in the lumbar but not thoracic spinal cord is reduced in Gars^ΔETAQ/+^ mice

We have previously shown that lower motor neuron cell bodies and nuclei are smaller in the lumbar spinal cord of *Gars*^*C201R/+*^ mice, but not in thoracic segments; this occurs without a reduction in motor neuron number (Sleigh et al., 2023). Dissecting the same spinal cord segments and staining for the motor neuron marker choline acetyltransferase (ChAT) (**Fig. 5A**), we replicated this lumbar-specific phenotype in *Gars*^Δ*ETAQ*/+^ mice. Consistent with peripheral neuropathy, there was no difference in lower motor neuron numbers in either lumbar or thoracic spinal cord (**Fig. 5B**), whereas motor neuron cell body area was selectively diminished in lumbar motor neurons, but not thoracic (**Fig. 5C**). These data are consistent with the DRG and spindle analyses, supporting that peripheral nerves innervating the hind-limbs of *Gars*^Δ*ETAQ*/+^ mice are preferentially impacted.

**Figure 5.**
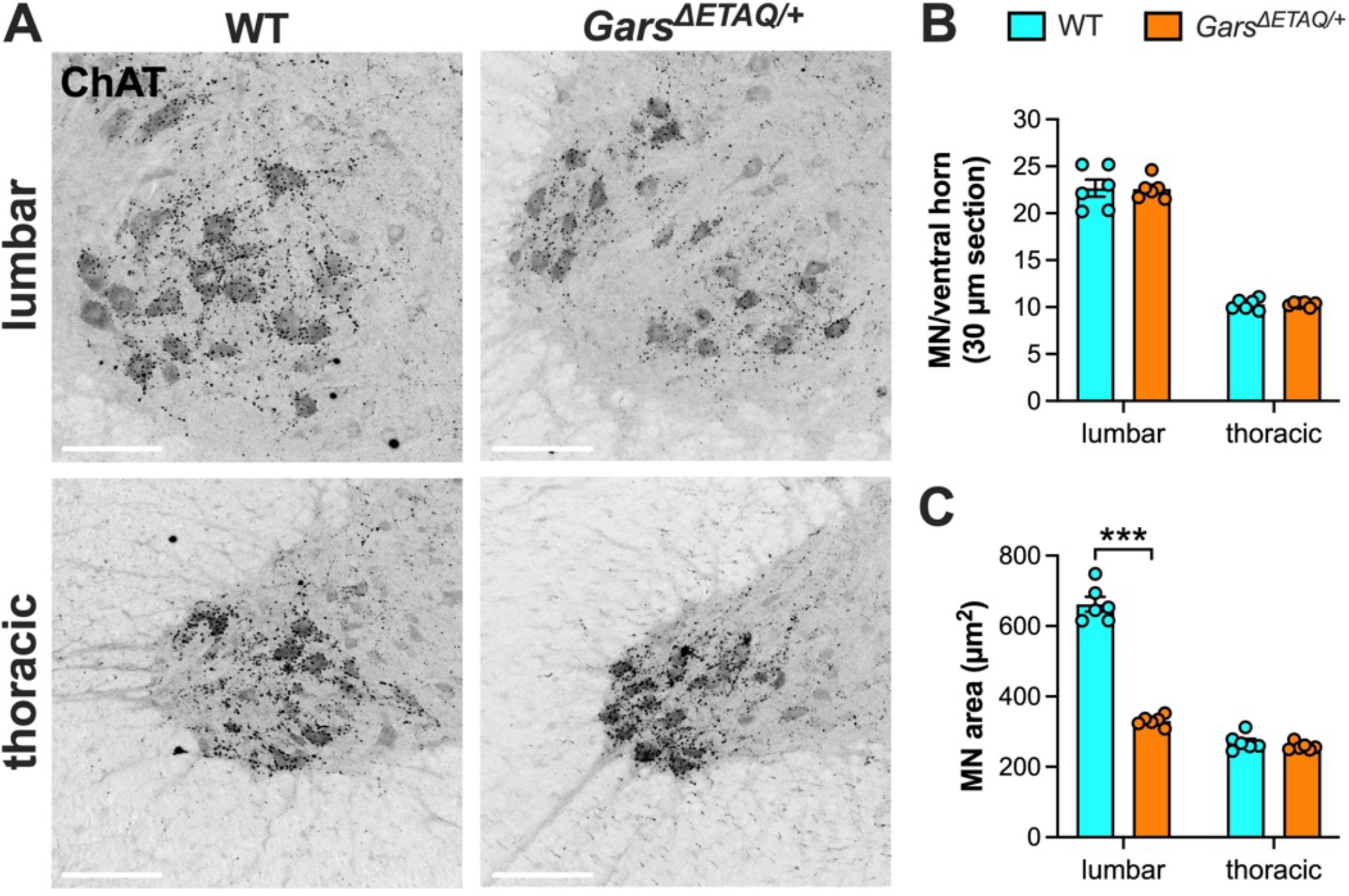
Lumbar motor neurons, but not thoracic motor neurons, display reduced areas in *Gars*^Δ*ETAQ*/+^ mice. (**A**) Representative collapsed z-stack images of wild-type and *Gars*^Δ*ETAQ*/+^ lumbar (top) and thoracic (bottom) spinal cord sections stained for ChAT. Scale bars = 100 µm. (**B**) There is no loss of motor neurons in the lumbar or thoracic spinal cord of *Gars*^Δ*ETAQ*/+^ mice (genotype *P* = 0.950, segment *P* < 0.001, interaction *P* = 0.925 two-way ANOVA). *Gars*^Δ*ETAQ*/+^ mice display reduced motor neuron areas in the lumbar spinal cord, but not the thoracic region (genotype *P* < 0.001, segment *P* < 0.001, interaction *P* < 0.001 two-way ANOVA). For all panels, data were generated from 3 month-old mice; \*\*\**P* < 0.001 Fisher’s post-hoc test; *n* = 6.

### 3.6 NMJ innervation and maturation are selectively impaired in Gars^ΔETAQ/+^mice

To further test the hypothesis that *Gars*^Δ*ETAQ*/+^ hind-limbs are more vulnerable to neuropathy, we examined the NMJ, an important pathological target in CMT (Snape and Sleigh, 2026). Indeed, the NMJ is selectively affected in mouse models for CMT2D (Seburn et al., 2006, Sleigh et al., 2014b, Spaulding et al., 2016, Sleigh et al., 2020b), with hind-limb muscles generally being most vulnerable to degeneration in *Gars*^*C201R/+*^ mutants (Sleigh et al., 2020b).

We therefore dissected some of the same muscles previously analysed in other CMT2D strains – the ETA of the upper fore-limb, the lumbricals of the fore-paw, and the FDB and lumbricals of the hind-paw – and stained them with fluorescent α-bungarotoxin (α-BTX) to visualise post-synaptic AChRs and antibodies against pan-synaptic vesicle 2 (SV2) and neurofilament (2H3) to label lower motor neurons (**Fig. 6A**). By evaluating the overlap between motor nerve terminals and AChRs, NMJs can be categorised as fully innervated, partially denervated or fully denervated, providing a quantitative measure of distal motor neuropathy.

**Figure 6.**
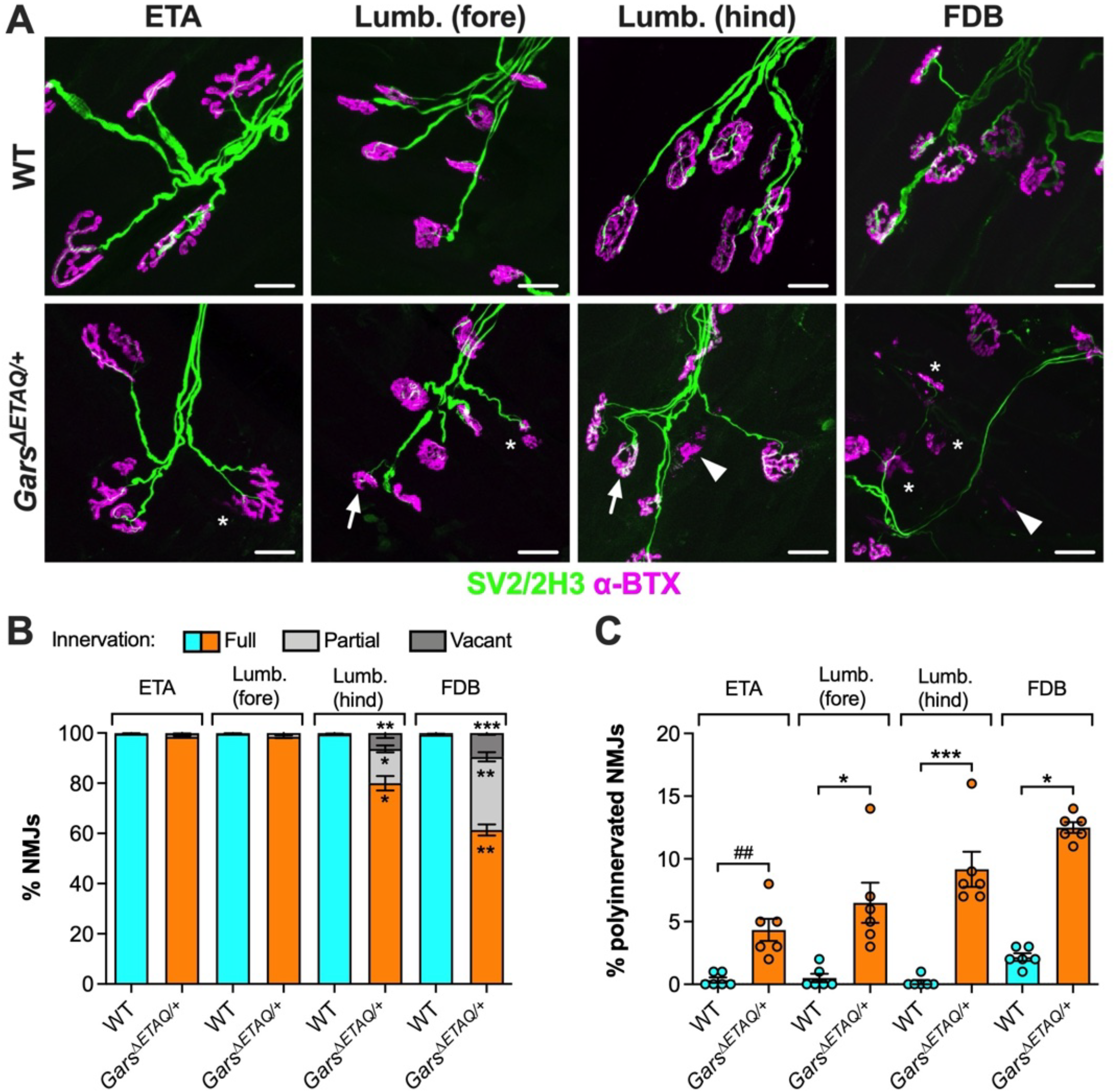
Neuromuscular junction innervation and maturation are more impaired in *Gars*^Δ*ETAQ*/+^ hind-limb than fore-limb muscles. (**A**) Representative collapsed z-stack confocal images of NMJs in ETA, fore-paw lumbrical [Lumb. (fore)], hind-paw lumbrical [Lumb. (hind)] and FDB muscles in wild-type and *Gars*^Δ*ETAQ*/+^ mice. Lower motor neurons are visualised using a combination of anti-SV2/anti-2H3 (green), and post-synaptic AChRs with α-BTX (magenta). Arrows highlight polyinnervated NMJs, asterisks partially denervated NMJs, and arrowheads fully denervated NMJs. Scale bars = 20 µm. (**B**) Muscles from *Gars*^Δ*ETAQ*/+^ mice display selective vulnerability to NMJ denervation: the ETA and fore-paw lumbricals show limited pathology, while hind-paw lumbrical and FDB muscles show marked denervation. (**C**) All assessed muscles show increased polyinnervation in *Gars*^Δ*ETAQ*/+^ mice, indicative of disturbed maturation. For all panels, data were generated from 3 month-old mice; genotypes were compared using Kruskall-Wallis tests; \**P* < 0.05, \*\**P* < 0.01, \*\*\**P* < 0.001 Dunn’s post-hoc test; ^##^*P* < 0.01 Mann-Whitney *U* test; *n* = 6.

Similar to *Gars*^*C201R/+*^ mice, we observed little to no pathology in the ETA and fore-limb lumbricals of *Gars*^Δ*ETAQ*/+^ mutants, whereas the hind-limb lumbricals and FDB muscles showed marked denervation (**Fig. 6B**). These differences in denervation were significant between all *Gars*^Δ*ETAQ*/+^ muscles, except the ETA and fore-limb lumbricals (Supplementary Figure 4A). Thus, there is a clear distal-to-proximal gradient of denervation across the evaluated *Gars*^Δ*ETAQ*/+^ muscles.

We have previously identified a correlation between NMJ denervation and synaptic maturity in both *Gars*^*Nmf249/+*^ and *Gars*^*C201R/+*^ mice, such that muscles with the greatest percentage of degenerated synapses also show the highest levels of polyinnervation (*i.e*., those with delayed synapse elimination) (Sleigh et al., 2014b, Sleigh et al., 2020b). Similarly, we found that *Gars*^Δ*ETAQ*/+^ muscles displayed a greater percentage of polyinnervation than wild-type (**Fig. 6C**). Polyinnervation was highest in the FDB of *Gars*^Δ*ETAQ*/+^ mice, which showed significantly greater levels than the ETA and fore-limb lumbricals (Supplementary Figure 4B). Correlating NMJ denervation (combining partial and full categories) with polyinnervation, we observed the same positive relationship as previously identified (Supplementary Figure 4C), indicating that the association between delayed synaptic maturation and denervation is a consistent feature of CMT2D mice.

### 3.7 In vivo axonal transport of signalling endosomes is selectively perturbed in Gars^ΔETAQ/+^ motor neurons

Impairments in axonal transport have been shown to precede axonal degeneration in many different models of inherited and acquired neuropathy (Prior et al., 2017), suggesting at least a contributory role in disease. Furthermore, we have previously identified that *in vivo* axonal transport of signalling endosomes becomes dysfunctional in motor axons of *Gars*^*C201R/+*^ mice between 2 weeks and 1 month of age, and is perturbed in *Gars*^Δ*ETAQ*/+^ mutants by 1 month (Sleigh et al., 2023)

To extend these findings, we injected the fluorescent probe H_C_T into the tibialis anterior, gastrocnemius or fore-paw of 3 month-old *Gars*^Δ*ETAQ*/+^ mice. The probe is taken up at the NMJ and loaded into retrogradely transported signalling endosomes, which can be time-lapse imaged and tracked in exposed peripheral nerves to determine their dynamic properties (**Fig. 7A**) (Tosolini et al., 2021, Lang et al., 2023). Hence, injecting different muscles and the fore-paw enables evaluation of axonal transport in motor neurons innervating muscles in distinct regions of the body. Indeed, the sciatic nerve was imaged following H_C_T injection into the tibialis anterior and gastrocnemius, whereas the median/ulnar nerves were imaged after injection into the fore-paw.

**Figure 7.**
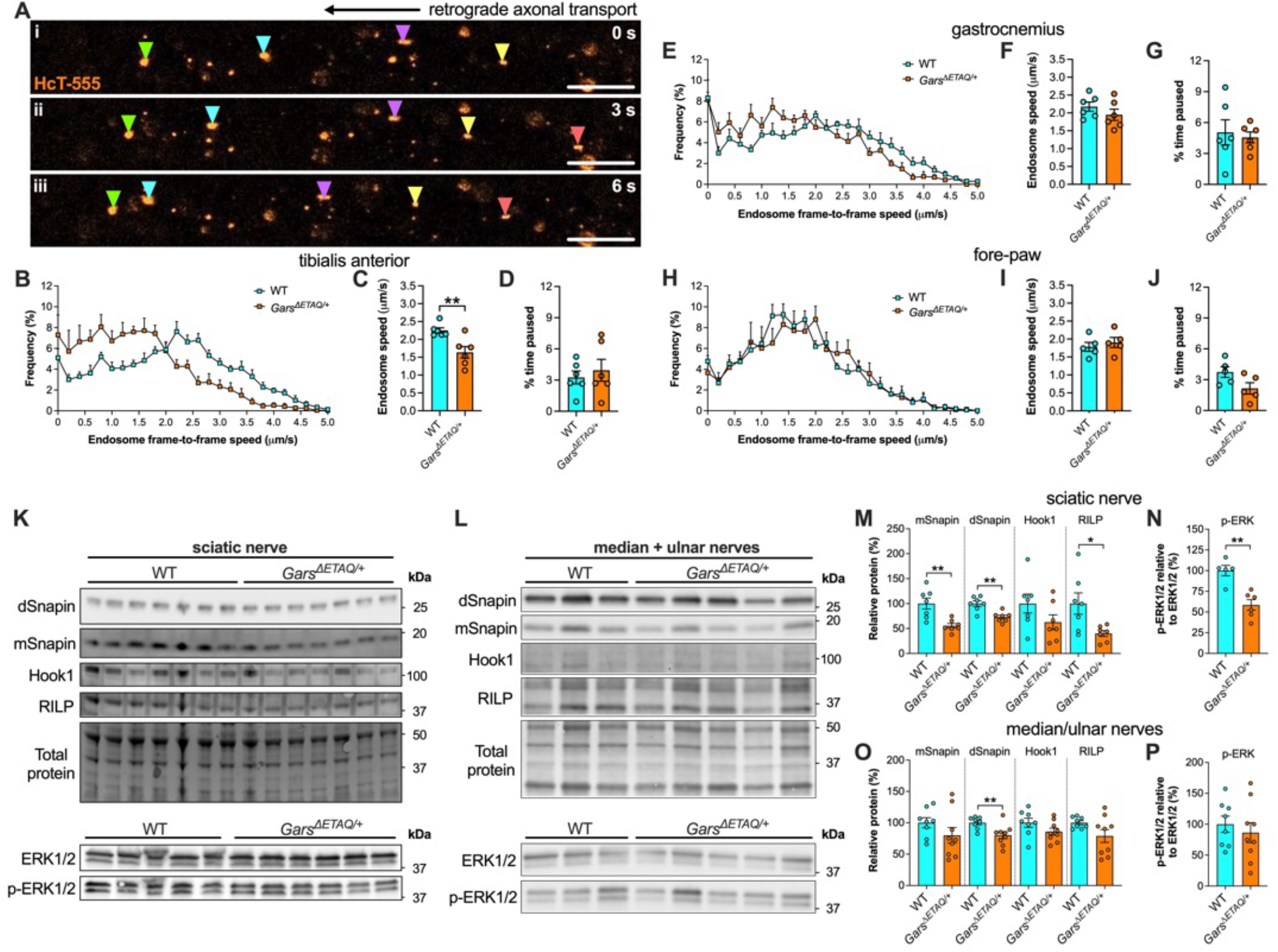
*In vivo* axonal transport of signaling endosomes is selectively disrupted in *Gars*^Δ*ETAQ*/+^ mice, linked with reduced endosome adaptor levels and impaired ERK1/2 phosphorylation. (**A**) Representative time-lapse confocal images of signalling endosomes labelled with an atoxic fluorescent fragment of tetanus neurotoxin (H_C_T-555, red) being retrogradely transported within peripheral nerve axons of live, anaesthetised mice. H_C_T-positive endosomes are individually tracked to quantitatively assess their dynamics. Colour-coded arrowheads identify five different endosomes. Scale bars = 10 µm. (**B**) Frame-to-frame speed histogram of signalling endosomes being transported within motor neurons innervating the tibialis anterior of wild-type and *Gars*^Δ*ETAQ*/+^ mice. (**C-D**) The speed of signalling endosome axonal transport is reduced in *Gars*^Δ*ETAQ*/+^ motor neurons innervating the tibialis anterior (C, *P* = 0.006), without a reduction in the percentage of time paused (D, *P* = 0.581). (**E**) Frame-to-frame speed histogram of signalling endosomes being transported within motor neurons innervating the gastrocnemius muscle of wild-type and *Gars*^Δ*ETAQ*/+^ mice. (**F-G**) There is no difference between genotypes in signalling endosome speed (F, *P* = 0.288) or percentage of time paused (G, *P* = 0.720) in motor neurons innervating the gastrocnemius. (**H**) Frame-to-frame speed histogram of signalling endosomes being transported within motor neurons innervating fore-paw muscles of wild-type and *Gars*^Δ*ETAQ*/+^ mice. (**I-J**) There is no difference between genotypes in signalling endosome speed (I, *P* = 0.531) or the percentage of time paused (J, *P* = 0.070) in motor neurons innervating the fore-paw muscles. (**K-L**) Western blots of dynein adaptor proteins Snapin, Hook1 and RILP (top), as well as total and phosphorylated ERK1/2 (bottom) in sciatic nerve (K) and median/ulnar nerve (L) lysates from wild-type and *Gars*^Δ*ETAQ*/+^ mice. (**M-N**) Densitometric analyses show reduced levels of monomeric Snapin (M, mSnapin, *P* = 0.003), dimeric Snapin (M, dSnapin, *P* = 0.001) and RILP (M, *P* = 0.019), as well as reduced ERK1/2 phosphorylation (N, *P* = 0.001) in *Gars*^Δ*ETAQ*/+^ sciatic nerves. Hook1 levels were not significantly affected (M, *P* = 0.146). (**O-P**) Densitometric analyses show that levels of mSnapin (O, *P* = 0.208), Hook1 (O, *P* = 0.134), RILP (O, *P* = 0.082) and phosphorylated ERK1/2 (P, *P* = 0.527) are unaffected in *Gars*^Δ*ETAQ*/+^ median/ulnar nerves. dSnapin levels were significantly reduced (O, *P* = 0.009). For all panels, data were generated from 3 month-old mice; genotypes were compared using unpaired *t*-tests; \**P* < 0.05, \*\**P* < 0.01; *n* = 5-6 (A-J), 5-7 (M-N) and 8-9 (O-P).

We observed a clear slowing of signalling endosomes in motor neurons innervating the tibialis anterior, without changes in the percentage of time spent paused (**Fig. 7B-D**). In contrast, motor neurons innervating the gastrocnemius and fore-paw muscles displayed no transport impairments (**Fig. 7E-J**). Together, these data confirm that *Gars*^Δ*ETAQ*/+^ mice display a selective disruption of *in vivo* signalling endosome axonal transport that correlates with the general pattern of neuropathy in this CMT2D model.

We have previously linked endosomal transport deficits in *Gars*^*C201R/+*^ mice to alterations in adaptor proteins required to connect dynein to signalling endosomes (Jordens et al., 2001, Cai et al., 2010, Olenick et al., 2019), and to reduced activation of Simkin, Paulo-Ramos *et al*., July 2026 – bioRxiv preprint ERK1/2, a key kinase that regulates signalling endosome transport initiation and speeds (Mitchell et al., 2012, Sleigh et al., 2023). To assess whether similar changes are observed in *Gars*^Δ*ETAQ*/+^ mice, we dissected sciatic nerves and combined median/ulnar nerves, and performed western blotting on their lysates. We identified reductions in monomeric snapin (mSnapin), dimeric snapin (dSnapin) and RILP levels in sciatic nerves (**Fig. 7K, M**), whereas only dSnapin was diminished in median/ulnar nerves (**Fig. L, O**). We also saw an impairment in ERK1/2 phosphorylation in sciatic nerves (**Fig. 7N**), but not in median/ulnar nerves (**Fig. 7P**).

Together, these data indicate that reduced availability of endosome adaptors and dampened ERK1/2 activation associate with, and thus perhaps contribute to, the disruption in signalling endosome axonal transport.

## Discussion

### 4.1 Peripheral nerve vulnerability and resistance

Peripheral nerves are not equally affected by neuropathy-causing mutations. As a consequence, CMT is often described as ‘length-dependent’, such that there is a priority for dysfunction in motor and sensory nerves innervating the feet, followed by the hands. However, mutations in a small subset of the more than 130 CMT genes, including *GARS1*, have been reported to drive an upper-limb predominant phenotype (Sivakumar et al., 2005, McMacken et al., 2021). In fact, neuropathy caused by *GARS1* mutations is often first observed in select intrinsic hand muscles (thenar and first dorsal interossei), with bilateral foot and peroneal muscle weakness emerging on average about three years later in CMT2D, but in only half of dHMNV patients (Sivakumar et al., 2005). The general pattern of neuropathy caused by *GARS1* mutations thus does not fit the ‘length-dependent’ model of CMT. In addition, the hypothenar muscles of the hand are relatively spared, despite having a motor pool that is similar in size and spinal cord location to that of the thenar muscles (Sivakumar et al., 2005). Hence, muscles innervated by axons of similar length can show differential vulnerability to CMT2D, as previously observed in *Gars*^*C201R/+*^ mice (Sleigh et al., 2020b), and now in *Gars*^Δ*ETAQ*/+^ mutants. This is reflected in the differential levels of NMJ denervation in hind-paw lumbricals and FDB muscles, as well as the selective axonal transport disruption in the tibialis anterior but not the gastrocnemius.

The cause for this selectivity remains unknown, but the pattern of degeneration can provide clues about disease pathogenesis. Indeed, we recently identified that distal denervation in *Gars*^*C201R/+*^ mice closely correlates with the extent of post-natal synaptic growth (Sleigh et al., 2020b) and availability of TrkB/BDNF in muscles (Sleigh et al., 2023), hinting that intrinsic features of the NMJ may contribute to selective motor neuron vulnerability. In the future, it will therefore be informative to evaluate AChR stability at the NMJ across CMT2D muscles and determine how TrkB/BDNF levels impact this.

Contrasting with the clinical onset, but matching the *Gars*^*C201R/+*^ phenotype (Sleigh et al., 2020a, Sleigh et al., 2020b), *Gars*^Δ*ETAQ*/+^ mice display clear vulnerability in motor and sensory nerves that innervate the hind-limbs, with relative sparing of those connecting to fore-limb muscles. The cause of the discrepancy between mice and humans remains undetermined. It is possible that the biomechanical context of quadrupedal walking rather than bipedally may protect fore-limb nerves from degeneration, or that the greater divergence in muscle and bone anatomy of the fore-limbs compared to the hind-limbs contributes to this effect (Ziermann et al., 2021). However, these explanations are unlikely, as mouse models of spinal muscular atrophy show the greatest pathology in proximal motor neurons (Woschitz et al., 2022). A few *GARS1* mutations (e.g., G598A) are reported to cause greater weakness in the feet and legs (James et al., 2006), so another possibility is that the mutations modelled in mice are simply lower-limb predominant; however, the relative severity across limbs was not reported in the young patient with the ΔETAQ mutation. Alternatively, fore-limb muscles other than those evaluated here may display pathology, akin to the difference between thenar and hypothenar muscles. Supporting this, it was recently shown that the proximal levator auris longus muscle in *Gars*^Δ*ETAQ*/+^ mice displays a modest reduction in NMJ innervation (Funke et al., 2026). Nonetheless, there is a much more striking loss of neuromuscular connectivity in hind-paw muscles across the mutant *Gars* alleles, consistent with a hind-limb predominance. It is conceivable that the molecular composition of cell bodies, axons and the NMJ (either at distal motor terminals, within the synaptic basal lamina, terminal Schwann cells or muscle) differ between mice and humans in a manner that makes peripheral nerves more or less susceptible to *GARS1* mutations. Hence, future studies should exploit the robust and selective degeneration of peripheral nerves in CMT2D mice to elucidate pathomechanisms and identify potential therapeutically targetable pathways.

### 4.2 Relative severity of sex and strain

Other than X-linked forms of neuropathy, the prevalence and severity of CMT are generally similar in females and males (Mladenovic et al., 2011, Theadom et al., 2019). Nevertheless, this is not the case for all subtypes – for example, in a cohort of 144 SORD neuropathy patients, the condition was more common (≈2:1) and trending to being more severe in males (Cortese et al., 2025), a finding that was replicated in a rat model of the disease (Rebelo et al., 2024). There is currently no evidence to support sex-specific differences in *GARS1*-related neuropathies, but it has not yet been systematically assessed, likely due to patient rarity, nor has it been rigorously evaluated in CMT2D mice. We therefore analysed several phenotypes using sufficient numbers of both females and males to determine whether there is any evidence for sex-specific differences in *Gars*^Δ*ETAQ*/+^ mice. Mutant males showed lower relative body weight compared to mutant females; however, no sex differences were observed in plasma NfL and periaxin levels, or sensory behaviour and motor function, suggesting that the severity of neuropathy is similar in female and male *Gars*^Δ*ETAQ*/+^ mutants. All other phenotypic assessments were consistent with this, albeit with smaller sample sizes and without statistical comparison (see **Supplementary Table 1**).

By three months, *Gars*^Δ*ETAQ*/+^ mice lose ≈10% of their axons within the motor branch of the femoral nerve, whereas *Gars*^*Nmf249/+*^ and *Gars*^*C201R/+*^ mutants show ≈30% and no loss, respectively (Seburn et al., 2006, Achilli et al., 2009, Morelli et al., 2019). Hence, *Gars*^Δ*ETAQ*/+^ mice are reported as having an intermediate phenotype; however, this has not been formally tested, nor have additional phenotypes been compared across mutants. In the present study, the only direct comparison between *Gars*^*C201R/+*^ and *Gars*^Δ*ETAQ*/+^ mice was the plasma periaxin measurements, which were higher in *Gars*^Δ*ETAQ*/+^ mutants but did not reach statistical significance (*P* = 0.079 Mann-Whitney *U* test). Comparing motor and sensory behaviour analyses in *Gars*^Δ*ETAQ*/+^ mice with those previously performed in 3 month-old *Gars*^*C201R/+*^ mutants (Sleigh et al., 2017), we observed a very similar extent of dysfunction across all tests, except for mechanical nociception. *Gars*^*C201R/+*^ mice displayed hypersensitivity to a noxious mechanical stimulus at both 1 and 3 months, whereas *Gars*^Δ*ETAQ*/+^ mutants showed a decreased response. As we observed similar numbers of peripherin+ neurons in the lumbar DRG across studies (*Gars*^Δ*ETAQ*/+^ 75.7 ± 1.0 % vs. *Gars*^*C201R/+*^ 75.9 ± 2.8 %; *P* = 0.848 unpaired *t*-test) (Sleigh et al., 2017), the distinction between strains is unlikely to reflect a proximal disruption in nociceptor number. Instead, the reduced mechanical nociception in *Gars*^Δ*ETAQ*/+^ mice is perhaps caused by a selective degeneration of nociceptor terminals in the skin. This remains to be confirmed, but is consistent with *Gars*^Δ*ETAQ*/+^ mice being more severely impacted than *Gars*^*C201R/+*^ mutants, as is the reduced size of their lumbar motor neurons across studies (330.3 ± 6.0 µm2 vs. 403.2 ± 12.3 µm2; *P* < 0.001 unpaired *t*-test) (Sleigh et al., 2023). That being said, we observed a similar decline in soleus muscle spindle number and innervation, but a lower percentage of denervated NMJs across *Gars*^Δ*ETAQ*/+^ versus *Gars*^*C201R/+*^ muscles (Sleigh et al., 2020b). This may be due to samples not being processed in parallel, but suggests that not all phenotypes are similar or more severe in *Gars*^Δ*ETAQ*/+^ mice. Alternatively, by evaluating the percentage of NMJ denervation without considering the total number of neuromuscular synapses, we may be underestimating the extent of pathology, since loss of neuromuscular connectivity leads to AChR dispersal (Comley et al., 2022). If a rapid wave of NMJ degeneration occurs well before 3 months in *Gars*^Δ*ETAQ*/+^ but not *Gars*^*C201R/+*^ muscles, the identified denervation percentages could belie the magnitude of pathology. Overall, our data support that the *Gars*^Δ*ETAQ*/+^ and *Gars*^*C201R/+*^ alleles are broadly similar in severity, with some elements of the *Gars*^Δ*ETAQ*/+^ phenotype being marginally worse; however, this requires confirmation through direct side-by-side assessments.

### 4.3 CMT2D biomarkers

Developing blood-based biomarkers for CMT that measure axonal damage and disease progression will be an important addition to patient outcome measures for assessing therapeutic interventions in clinical trials (Rossor et al., 2020). Similarly, in combination with phenotypic assessments, biomarkers in mice can be used to objectively test pre-clinical therapeutics. Surprisingly, despite its systematic release upon axon degeneration, NfL levels were lower in *Gars*^Δ*ETAQ*/+^ plasma compared to wild-type. This phenomenon has also been reported to intermittently occur in *Gars*^*C201R/+*^ mice, but not *Gars*^*Nmf249/+*^ (Rossor et al., 2022). This may reflect activation of the integrated stress response impairing the production of NfL (Spaulding et al., 2021), and/or diminished slow axonal transport of NfL towards peripheral nerve terminals (Trivedi et al., 2007, Kotaich et al., 2023). Consistent with these ideas, both *Gars*^Δ*ETAQ*/+^ and *Gars*^*C201R/+*^ mice display a reduction in peripheral axon calibres (Achilli et al., 2009, Morelli et al., 2019), which is dependent on neurofilaments (Hoffman et al., 1987). Alternatively, mutant GlyRS could impact the stability of NfL mRNA, as has been shown for mutant SOD1-linked to amyotrophic lateral sclerosis (Ge et al., 2005). In contrast to NfL, we saw no significant changes between genotypes in another intermediate filament protein, peripherin; this may reflect differences in protein half-life or assay sensitivity, but further timepoints should be assessed before ruling out peripherin as a useful biomarker.

We also observed a clear increase in levels of plasma periaxin, which is exclusively expressed in Schwann cells rather than neurons and was recently identified as a biomarker for demyelination (Bellanti et al., 2025). This fits with a Schwann cell disruption in *Gars*^Δ*ETAQ*/+^ mice, which is not clinically observed in CMT2D patients, as the disease is classified as an axonal neuropathy. Nevertheless, it is consistent with the *Gars*^Δ*ETAQ*/+^ mice displaying a decline in NCV and reduced myelin thickness (Morelli et al., 2019, Ozes et al., 2021).

Counter to observations in *Gars*^*Nmf249/+*^ and *Gars*^*C201R/+*^ mice (Jennings et al., 2022), we did not detect a change in circulating levels of the cytokine Gdf15 within the blood of *Gars*^Δ*ETAQ*/+^ mutants. This may reflect an allele-specific difference; however, given that increased Gdf15 has been reported across multiple CMT subtypes and in corresponding mouse models, this is unlikely. Alternatively, the difference may relate to disease timepoint (5 months vs. 3 months here) or the blood fraction analysed (serum vs. plasma here).

### 4.4 Limitations

There are several limiting elements to the present study. Firstly, we assessed *Gars*^Δ*ETAQ*/+^ mice at a single timepoint and therefore do not know when the identified phenotypes first manifest or whether they progressively worsen. Understanding the temporal profile of disease is important for determining the pre-symptomatic window, and for determining whether pathology sufficiently increases with age to enable post-symptomatic therapy testing; this is particularly relevant given the developmental phenotypes observed in CMT2D mice (He et al., 2015, Sleigh et al., 2017). Secondly, several experiments were not sufficiently powered to detect potential sex differences in neuropathology, although this is perhaps unlikely given the similarity in sensory and motor behavioural deficits between sexes. Finally, assessment of NMJ neurotransmission in addition to morphology is important to fully determine peripheral nerve dysfunction, as structurally normal synapses can be functionally impaired (Snape and Sleigh, 2026).

### 4.5 Conclusions

We show that *Gars*^Δ*ETAQ*/+^ mice display robust and selective peripheral nerve pathology, underlying clear functional deficits. Moreover, we confirmed that NfL and periaxin are useful blood biomarkers for objective assessment of neurodegeneration in this model, that the sensory nervous system displays clear functional and morphological disruption, and that there is a similar pattern of NMJ denervation as that observed in *Gars*^*C201R/+*^ mice. This work not only confirms the usefulness of *Gars*^Δ*ETAQ*/+^ mice for testing treatments and evaluating mechanisms underlying selective nerve vulnerability, but it also demonstrates the validity of the *Gars*^*Nmf249/+*^ and *Gars*^*C201R/+*^ strains. While the respective spontaneous and mutagen-induced mutations may never be identified in humans, these models display a strikingly similar phenotype to *Gars*^Δ*ETAQ*/+^ mice, confirming that disease model mice do not need to harbour a patient-sourced mutation in order to reproduce key features of a disease.

## Acknowledgements

We thank Prof. Giampietro Schiavo (UCL) for support and encouragement of the work. This project was funded by Medical Research Council awards MR/S006990/1 and MR/Y010949/1 (JNS); a Rosetrees Trust grant M806 (JNS); the UCL Neurogenetic Therapies Programme funded by The Sigrid Rausing Trust (JNS); the UCL Therapeutic Acceleration Support scheme supported by funding from MRC IAA 2021 UCL MR/X502984/1 (JNS); a Muscular Dystrophy UK grant 24GRO-PG24-0719-1 (JNS), a Human Frontier Science Program Long-Term Fellowship LT000220/2017-L (SS); and a National Institutes of Health award R37 NS05415 (RWB). HZ is a Wallenberg Scholar and a Distinguished Professor at the Swedish Research Council supported by grants from the Swedish Research Council (#2023-00356, #2022-01018 and #2019-02397), the European Union’s Horizon Europe research and innovation programme under grant agreement No 101053962, Swedish State Support for Clinical Research (#ALFGBG-71320), the Alzheimer Drug Discovery Foundation (ADDF), USA (#201809-2016862), the AD Strategic Fund and the Alzheimer’s Association (#ADSF-21-831376-C, #ADSF-21-831381-C, #ADSF-21-831377-C, and #ADSF-24-1284328-C), the European Partnership on Metrology, co-financed from the European Union’s Horizon Europe Research and Innovation Programme and by the Participating States (NEuroBioStand, #22HLT07), the Bluefield Project, Cure Alzheimer’s Fund, the Olav Thon Foundation, the Erling-Persson Family Foundation, Familjen Rönströms Stiftelse, Familjen Beiglers Stiftelse, Stiftelsen för Gamla Tjänarinnor, Hjärnfonden, Sweden (#FO2022-0270), the European Union’s Horizon 2020 research and innovation programme under the Marie Skłodowska-Curie grant agreement No 860197 (MIRIADE), the European Union Joint Programme – Neurodegenerative Disease Research (JPND2021-00694), the National Institute for Health and Care Research University College London Hospitals Biomedical Research Centre, the UK Dementia Research Institute at UCL (UKDRI-1003), and an anonymous donor. The graphical abstract was produced using www.biorender.com. Within the graphical abstract, Claude.ai was used to create the muscle spindle graphic and provide the green-to-red colour gradient of the nervous system.

## Author Contributions

Conceptualisation: JNS. Investigation: RLS, AP-R, QL, ERR, SS, DV-C, SL, RB, EV, VD, OS, JNS. Resources: RWB. Supervision: AH, HZ, MPL, JNS. Writing – original draft: JNS. Writing – review and editing: RLS, AP-R, DV-C, HZ, JNS. Funding acquisition: RWB, JNS.

## Conflict of Interest

HZ has served at scientific advisory boards and/or as a consultant for Abbvie, Acumen, Alamar, Alector, Alzinova, ALZpath, Amylyx, Annexon, Apellis, Artery Therapeutics, AZTherapies, Bioventix, Cognitact, Cognito Therapeutics, CogRx, Denali, Eisai, Enigma, Johnson & Johnson, LabCorp, Merck Sharp & Dohme, Merry Life, Nervgen, New Amsterdam, Novo Nordisk, Optoceutics, Passage Bio, Pinteon Therapeutics, Prothena, Quanterix, Red Abbey Labs, reMYND, Roche, Samumed, ScandiBio Therapeutics AB, Siemens Healthineers, Triplet Therapeutics, and Wave, has given lectures sponsored by Alzecure, BioArctic, Biogen, Cellectricon, Fujirebio, LabCorp, Lilly, Novo Nordisk, Oy Medix Biochemica AB, Roche, and WebMD, is a co-founder of Brain Biomarker Solutions in Gothenburg AB (BBS), which is a part of the GU Ventures Incubator Program, and is a shareholder of CERimmune Therapeutics (outside submitted work). The other authors declare no competing interests.

## References

Achilli, F., Bros-Facer, V., Williams, H.P., Banks, G.T., AlQatari, M., Chia, R., Tucci, V., Groves, M., Nickols, C.D., Seburn, K.L., Kendall, R., Cader, M.Z., Talbot, K., van Minnen, J., Burgess, R.W., Brandner, S., Martin, J.E., Koltzenburg, M., Greensmith, L., Nolan, P.M., Fisher, E.M., 2009. An ENU-induced mutation in mouse glycyl-tRNA synthetase (GARS) causes peripheral sensory and motor phenotypes creating a model of Charcot-Marie-Tooth type 2D peripheral neuropathy. Dis. Model. Mech. 2, 359–373. 10.1242/dmm.002527.

Antonellis, A., Ellsworth, R.E., Sambuughin, N., Puls, I., Abel, A., Lee-Lin, S.Q., Jordanova, A., Kremensky, I., Christodoulou, K., Middleton, L.T., Sivakumar, K., Ionasescu, V., Funalot, B., Vance, J.M., Goldfarb, L.G., Fischbeck, K.H., Green, E.D., 2003. Glycyl tRNA synthetase mutations in Charcot-Marie-Tooth disease type 2D and distal spinal muscular atrophy type V. Am. J. Hum. Genet. 72, 1293–1299. 10.1086/375039.

Bellanti, R., Keh, R.Y.S., Keddie, S., Chou, M.K.L., Misheva, M., Smyth, D., Baskozos, G., Moodley, K., Hart, M.S., Davies, A.J., Reilly, M.M., Rinaldi, S., Lunn, M.P., 2025. Plasma periaxin is a biomarker of peripheral nerve demyelination. Brain. 148, 4448–4460. 10.1093/brain/awaf234.

Benoy, V., Van Helleputte, L., Prior, R., d’Ydewalle, C., Haeck, W., Geens, N., Scheveneels, W., Schevenels, B., Cader, M.Z., Talbot, K., Kozikowski, A.P., Vanden Berghe, P., Van Damme, P., Robberecht, W., Van Den Bosch, L., 2018. HDAC6 is a therapeutic target in mutant GARS-induced Charcot-Marie-Tooth disease. Brain. 141, 673–687. 10.1093/brain/awx375.

Bonin, R.P., Bories, C., De Koninck, Y., 2014. A simplified up-down method (SUDO) for measuring mechanical nociception in rodents using von Frey filaments. Mol. Pain. 10, 26. 10.1186/1744-8069-10-26.

Bosco, L., Falzone, Y.M., Previtali, S.C., 2021. Animal models as a tool to design therapeutical strategies for CMT-like hereditary neuropathies. Brain Sci. 11, 1237. 10.3390/brainsci11091237.

Burgess, R.W., Storkebaum, E., 2023. tRNA dysregulation in neurodevelopmental and neurodegenerative diseases. Annu. Rev. Cell Dev. Biol. 39, 223–252. 10.1146/annurev-cellbio-021623-124009.

Burns, J., Timmerman, V., Laurà, M., Yiu, E.M., D’Antonio, M., Mukherjee-Clavin, B., De Winter, J., Scherer, S.S., 2026. Charcot-Marie-Tooth disease and related neuropathies. Nat. Rev. Dis. Primers. 12, 3. 10.1038/s41572-025-00679-2.

Cai, Q., Lu, L., Tian, J.H., Zhu, Y.B., Qiao, H., Sheng, Z.H., 2010. Snapin-regulated late endosomal transport is critical for efficient autophagy-lysosomal function in neurons. Neuron. 68, 73–86. 10.1016/j.neuron.2010.09.022.

Carter, R.J., Morton, J., Dunnett, S.B., 2001. Motor coordination and balance in rodents. Curr. Protoc. Neurosci. Chapter 8, Unit 8.12. 10.1002/0471142301.ns0812s15.

Comley, L.H., Kline, R.A., Thomson, A.K., Woschitz, V., Landeros, E.V., Osman, E.Y., Lorson, C.L., Murray, L.M., 2022. Motor unit recovery following Smn restoration in mouse models of spinal muscular atrophy. Hum. Mol. Genet. 31, 3107–3119. 10.1093/hmg/ddac097.

Cortese, A., Dohrn, M.F., Currò, R., Negri, S., Lassuthova, P., Pisciotta, C., Tozza, S., Al-Ajmi, A., Feng, C., Tomaselli, P.J., Fernandez-Eulate, G., Haddad, S., Laurà, M., Rossor, A.M., Vegezzi, E., Facchini, S., Sleigh, J.N., Rebelo, A., Beijer, D., Raposo, J., Saporta, M., Lauerova, B., Pernice, H.F., Achenbach, P., Schöne, U., Alon, T., Deschauer, M., Cordts, I., Obermaier, C.D., Winter, N., Creigh, P.D., Sowden, J.E., Rehbein, T., Magri, S., Bertini, A., Saveri, P., Ripellino, P., Huang, J., Nadaj-Pakleza, A., Ross, A., Holt, J.K.L., Brennan, K.M., Sukenik-Halevy, R., Bizaoui, V., Parman, Y., Battaloglu, E., Çakar, A., Alrohaif, H., Hammans, S., Kumar, K.R., Kennerson, M.L., Kayserili, H., Amado, D.A., Hahn, K., Valentino, P., Cavalcanti, F., Gaetano, C., Taroni, F., Braathen, G.J., Houlden, H., Stojkovic, T., Peric, S., Bolino, A., Previtali, S.C., Yi-Chung, L., Başak, A.N., Hamed, S.A., Rojas-García, R., Claeys, K.G., Marques, W., Sevilla, T., Schlotter-Weigel, B., Manganelli, F., Zhang, R., Herrmann, D.N., Scherer, S.S., Seeman, P., Pareyson, D., Reilly, M.M., Shy, M.E., Züchner, S., 2025. Genotype and phenotype spectrum of Charcot-Marie-Tooth disease due to mutations in *SORD*. Brain. 148, 3737–3747. 10.1093/brain/awaf021.

Deenen, J.C., Verbeek, A.L., Verschuuren, J.J., van Engelen, B.G., Voermans, N.C., 2025. Prevalence and incidence rates of 17 neuromuscular disorders: An updated review of the literature. J. Neuromuscul. Dis. 12, 713–722. 10.1177/22143602241313118.

Funke, J.R., Martinez, C., Pratt, S.L., Rice, A.D., Tadenev, A.L.D., Burgess, R.W., 2026. Neuromuscular junction dysfunction in a subset of Charcot-Marie-Tooth and related peripheral neuropathies mouse models. Neurobiol. Dis. 225, 107455. 10.1016/j.nbd.2026.107455.

Ge, W.W., Wen, W., Strong, W., Leystra-Lantz, C., Strong, M.J., 2005. Mutant copper-zinc superoxide dismutase binds to and destabilizes human low molecular weight neurofilament mRNA. J. Biol. Chem. 280, 118–124. 10.1074/jbc.M405065200.

Gibbs, K.L., Kalmar, B., Sleigh, J.N., Greensmith, L., Schiavo, G., 2016. *In vivo* imaging of axonal transport in murine motor and sensory neurons. J. Neurosci. Methods. 257, 26–33. 10.1016/j.jneumeth.2015.09.018.

Hargreaves, K., Dubner, R., Brown, F., Flores, C., Joris, J., 1988. A new and sensitive method for measuring thermal nociception in cutaneous hyperalgesia. Pain. 32, 77–88. 10.1016/0304-3959(88)90026-7.

He, W., Bai, G., Zhou, H., Wei, N., White, N.M., Lauer, J., Liu, H., Shi, Y., Dumitru, C.D., Lettieri, K., Shubayev, V., Jordanova, A., Guergueltcheva, V., Griffin, P.R., Burgess, R.W., Pfaff, S.L., Yang, X.L., 2015. CMT2D neuropathy is linked to the neomorphic binding activity of glycyl-tRNA synthetase. Nature. 526, 710–714. 10.1038/nature15510.

He, W., Zhang, H.M., Chong, Y.E., Guo, M., Marshall, A.G., Yang, X.L., 2011. Dispersed disease-causing neomorphic mutations on a single protein promote the same localized conformational opening. Proc. Natl. Acad. Sci. U. S. A. 108, 12307–12312. 10.1073/pnas.1104293108.

Hoffman, P.N., Cleveland, D.W., Griffin, J.W., Landes, P.W., Cowan, N.J., Price, D.L., 1987. Neurofilament gene expression: a major determinant of axonal caliber. Proc. Natl. Acad. Sci. U. S. A. 84, 3472–3476. 10.1073/pnas.84.10.3472.

James, P.A., Cader, M.Z., Muntoni, F., Childs, A.M., Crow, Y.J., Talbot, K., 2006. Severe childhood SMA and axonal CMT due to anticodon binding domain mutations in the *GARS* gene. Neurology. 67, 1710–1712. 10.1212/01.wnl.0000242619.52335.bc.

Jennings, M.J., Kagiava, A., Vendredy, L., Spaulding, E.L., Stavrou, M., Hathazi, D., Grüneboom, A., De Winter, V., Gess, B., Schara, U., Pogoryelova, O., Lochmüller, H., Borchers, C.H., Roos, A., Burgess, R.W., Timmerman, V., Kleopa, K.A., Horvath, R., 2022. NCAM1 and GDF15 are biomarkers of Charcot-Marie-Tooth disease in patients and mice. Brain. 145, 3999–4015. 10.1093/brain/awac055.

Jordens, I., Fernandez-Borja, M., Marsman, M., Dusseljee, S., Janssen, L., Calafat, J., Janssen, H., Wubbolts, R., Neefjes, J., 2001. The Rab7 effector protein RILP controls lysosomal transport by inducing the recruitment of dynein-dynactin motors. Curr. Biol. 11, 1680–1685. 10.1016/s0960-9822(01)00531-0.

Juneja, M., Burns, J., Saporta, M.A., Timmerman, V., 2019. Challenges in modelling the Charcot-Marie-Tooth neuropathies for therapy development. J. Neurol. Neurosurg. Psychiatry. 90, 58–67. 10.1136/jnnp-2018-318834.

Kalotay, E., Klugmann, M., Housley, G.D., Fröhlich, D., 2023. Dominant aminoacyl-tRNA synthetase disorders: lessons learned from *in vivo* disease models. Front. Neurosci. 17, 1182845. 10.3389/fnins.2023.1182845.

Keddie, S., Smyth, D., Keh, R.Y.S., Chou, M.K.L., Grant, D., Surana, S., Heslegrave, A., Zetterberg, H., Wieske, L., Michael, M., Eftimov, F., Bellanti, R., Rinaldi, S., Hart, M.S., Petzold, A., Lunn, M.P., 2023. Peripherin is a biomarker of axonal damage in peripheral nervous system disease. Brain. 146, 4562–4573. 10.1093/brain/awad234.

Kotaich, F., Caillol, D., Bomont, P., 2023. Neurofilaments in health and Charcot-Marie-Tooth disease. Front. Cell Dev. Biol. 11, 1275155. 10.3389/fcell.2023.1275155.

Lang, Q., Schiavo, G., Sleigh, J.N., 2023. *In vivo* imaging of axonal transport in peripheral nerves of rodent forelimbs. Neuronal Signal. 7, NS20220098. 10.1042/NS20220098.

McMacken, G., Whittaker, R.G., Charlton, R., Barresi, R., Lochmüller, H., Horvath, R., 2021. Inherited neuropathies with predominant upper limb involvement: genetic heterogeneity and overlapping pathologies. Eur. J. Neurol. 28, 297–304. 10.1111/ene.14514.

Mech, A.M., Brown, A.L., Schiavo, G., Sleigh, J.N., 2020. Morphological variability is greater at developing than mature mouse neuromuscular junctions. J. Anat. 237, 603–617. 10.1111/joa.13228.

Millere, E., Rots, D., Simren, J., Ashton, N.J., Kupats, E., Micule, I., Priedite, V., Kurjane, N., Blennow, K., Gailite, L., Zetterberg, H., Kenina, V., 2021. Plasma neurofilament light chain as a potential biomarker in Charcot-Marie-Tooth disease. Eur. J. Neurol. 28, 974–981. 10.1111/ene.14689.

Mitchell, D.J., Blasier, K.R., Jeffery, E.D., Ross, M.W., Pullikuth, A.K., Suo, D., Park, J., Smiley, W.R., Lo, K.W., Shabanowitz, J., Deppmann, C.D., Trinidad, J.C., Hunt, D.F., Catling, A.D., Pfister, K.K., 2012. Trk activation of the ERK1/2 kinase pathway stimulates intermediate chain phosphorylation and recruits cytoplasmic dynein to signaling endosomes for retrograde axonal transport. J. Neurosci. 32, 15495–15510. 10.1523/JNEUROSCI.5599-11.2012.

Mladenovic, J., Milic Rasic, V., Keckarevic Markovic, M., Romac, S., Todorovic, S., Rakocevic Stojanovic, V., Kisic Tepavcevic, D., Hofman, A., Pekmezovic, T., 2011. Epidemiology of Charcot-Marie-Tooth disease in the population of Belgrade, Serbia. Neuroepidemiology. 36, 177–182. 10.1159/000327029.

Mo, Z., Zhao, X., Liu, H., Hu, Q., Chen, X.Q., Pham, J., Wei, N., Liu, Z., Zhou, J., Burgess, R.W., Pfaff, S.L., Caskey, C.T., Wu, C., Bai, G., Yang, X.L., 2018. Aberrant GlyRS-HDAC6 interaction linked to axonal transport deficits in Charcot-Marie-Tooth neuropathy. Nat. Commun. 9, 1007. 10.1038/s41467-018-03461-z.

Montaño, J.A., Pérez-Piñera, P., García-Suárez, O., Cobo, J., Vega, J.A., 2010. Development and neuronal dependence of cutaneous sensory nerve formations: Lessons from neurotrophins. Microsc. Res. Tech. 73, 513–529. 10.1002/jemt.20790.

Morelli, K.H., Griffin, L.B., Pyne, N.K., Wallace, L.M., Fowler, A.M., Oprescu, S.N., Takase, R., Wei, N., Meyer-Schuman, R., Mellacheruvu, D., Kitzman, J.O., Kocen, S.G., Hines, T.J., Spaulding, E.L., Lupski, J.R., Nesvizhskii, A., Mancias, P., Butler, I.J., Yang, X.L., Hou, Y.M., Antonellis, A., Harper, S.Q., Burgess, R.W., 2019. Allele-specific RNA interference prevents neuropathy in Charcot-Marie-Tooth disease type 2D mouse models. J. Clin. Invest. 129, 5568–5583. 10.1172/JCI130600.

Motley, W.W., Seburn, K.L., Nawaz, M.H., Miers, K.E., Cheng, J., Antonellis, A., Green, E.D., Talbot, K., Yang, X.L., Fischbeck, K.H., Burgess, R.W., 2011. Charcot-Marie-Tooth-linked mutant GARS is toxic to peripheral neurons independent of wild-type GARS levels. PLoS Genet. 7, e1002399. 10.1371/journal.pgen.1002399.

Olenick, M.A., Dominguez, R., Holzbaur, E.L.F., 2019. Dynein activator Hook1 is required for trafficking of BDNF-signaling endosomes in neurons. J. Cell Biol. 218, 220–233. 10.1083/jcb.201805016.

Ozes, B., Moss, K., Myers, M., Ridgley, A., Chen, L., Murrey, D., Sahenk, Z., 2021. AAV1.NT-3 gene therapy in a CMT2D model: phenotypic improvements in GarsP278KY/+ mice. Brain Commun. 3, fcab252. 10.1093/braincomms/fcab252.

Pipis, M., Rossor, A.M., Laurà, M., Reilly, M.M., 2019. Next-generation sequencing in Charcot-Marie-Tooth disease: opportunities and challenges. Nat. Rev. Neurol. 15, 644–656. 10.1038/s41582-019-0254-5.

Prior, R., Van Helleputte, L., Benoy, V., Van Den Bosch, L., 2017. Defective axonal transport: A common pathological mechanism in inherited and acquired peripheral neuropathies. Neurobiol. Dis. 105, 300–320. 10.1016/j.nbd.2017.02.009.

Rebelo, A.P., Abad, C., Dohrn, M.F., Li, J.J., Tieu, E.K., Medina, J., Yanick, C., Huang, J., Zotter, B., Young, J.I., Saporta, M., Scherer, S.S., Walz, K., Züchner, S., 2024. SORD-deficient rats develop a motor-predominant peripheral neuropathy unveiling novel pathophysiological insights. Brain. 147, 3131–3143. 10.1093/brain/awae079.

Record, C.J., Pipis, M., Skorupinska, M., Blake, J., Poh, R., Polke, J.M., Eggleton, K., Nanji, T., Züchner, S., Cortese, A., Houlden, H., Rossor, A.M., Laurà, M., Reilly, M.M., 2024. Whole genome sequencing increases the diagnostic rate in Charcot-Marie-Tooth disease. Brain. 147, 3144–3156. 10.1093/brain/awae064.

Reilly, M.M., Murphy, S.M., Laurà, M., 2011. Charcot-Marie-Tooth disease. J. Peripher. Nerv. Syst. 16, 1–14. 10.1111/j.1529-8027.2011.00324.x.

Rhymes, E.R., Simkin, R.L., Qu, J., Villarroel-Campos, D., Surana, S., Tong, Y., Shapiro, R., Burgess, R.W., Yang, X.L., Schiavo, G., Sleigh, J.N., 2024. Boosting BDNF in muscle rescues impaired axonal transport in a mouse model of DI-CMTC peripheral neuropathy. Neurobiol. Dis. 195, 106501. 10.1016/j.nbd.2024.106501.

Rice, A.D., Tadenev, A.L.D., Hines, T.J., Funke, J.R., Burgess, R.W., 2025. SARM1 inhibition in three mouse models of Charcot-Marie-Tooth disease. J. Peripher. Nerv. Syst. 30, e70053. 10.1111/jns.70053.

Rohrer, J.D., Woollacott, I.O., Dick, K.M., Brotherhood, E., Gordon, E., Fellows, A., Toombs, J., Druyeh, R., Cardoso, M.J., Ourselin, S., Nicholas, J.M., Norgren, N., Mead, S., Andreasson, U., Blennow, K., Schott, J.M., Fox, N.C., Warren, J.D., Zetterberg, H., 2016. Serum neurofilament light chain protein is a measure of disease intensity in frontotemporal dementia. Neurology. 87, 1329–1336. 10.1212/WNL.0000000000003154.

Rossor, A.M., Kapoor, M., Wellington, H., Spaulding, E., Sleigh, J.N., Burgess, R.W., Laurà, M., Zetterberg, H., Bacha, A., Wu, X., Heslegrave, A., Shy, M.E., Reilly, M.M., 2022. A longitudinal and cross-sectional study of plasma neurofilament light chain concentration in Charcot-Marie-Tooth disease. J. Peripher. Nerv. Syst. 27, 50–57. 10.1111/jns.12477.

Rossor, A.M., Shy, M.E., Reilly, M.M., 2020. Are we prepared for clinical trials in Charcot-Marie-Tooth disease? Brain Res. 1729, 146625. 10.1016/j.brainres.2019.146625.

Seburn, K.L., Nangle, L.A., Cox, G.A., Schimmel, P., Burgess, R.W., 2006. An active dominant mutation of glycyl-tRNA synthetase causes neuropathy in a Charcot-Marie-Tooth 2D mouse model. Neuron. 51, 715–726. 10.1016/j.neuron.2006.08.027.

Simkin, R.L., Rhymes, E.R., Lang, Q., Birsa, N., Sleigh, J.N., 2025. Dissection and whole-mount immunofluorescent staining of mouse hind paw muscles for neuromuscular junction analysis. Bio Protoc. 15, e5315. 10.21769/BioProtoc.5315.

Sivakumar, K., Kyriakides, T., Puls, I., Nicholson, G.A., Funalot, B., Antonellis, A., Sambuughin, N., Christodoulou, K., Beggs, J.L., Zamba-Papanicolaou, E., Ionasescu, V., Dalakas, M.C., Green, E.D., Fischbeck, K.H., Goldfarb, L.G., 2005. Phenotypic spectrum of disorders associated with glycyl-tRNA synthetase mutations. Brain. 128, 2304–2314. 10.1093/brain/awh590.

Sleigh, J.N., Burgess, R.W., Gillingwater, T.H., Cader, M.Z., 2014a. Morphological analysis of neuromuscular junction development and degeneration in rodent lumbrical muscles. J. Neurosci. Methods. 227, 159–165. 10.1016/j.jneumeth.2014.02.005.

Sleigh, J.N., Dawes, J.M., West, S.J., Wei, N., Spaulding, E.L., Gómez-Martín, A., Zhang, Q., Burgess, R.W., Cader, M.Z., Talbot, K., Yang, X.L., Bennett, D.L., Schiavo, G., 2017. Trk receptor signaling and sensory neuron fate are perturbed in human neuropathy caused by *Gars* mutations. Proc. Natl. Acad. Sci. U. S. A. 114, E3324–E3333. 10.1073/pnas.1614557114.

Sleigh, J.N., Grice, S.J., Burgess, R.W., Talbot, K., Cader, M.Z., 2014b. Neuromuscular junction maturation defects precede impaired lower motor neuron connectivity in Charcot-Marie-Tooth type 2D mice. Hum. Mol. Genet. 23, 2639–2650. 10.1093/hmg/ddt659.

Sleigh, J.N., Mech, A.M., Aktar, T., Zhang, Y., Schiavo, G., 2020a. Altered sensory neuron development in CMT2D mice is site-specific and linked to increased GlyRS levels. Front. Cell. Neurosci. 14, 232. 10.3389/fncel.2020.00232.

Sleigh, J.N., Mech, A.M., Schiavo, G., 2020b. Developmental demands contribute to early neuromuscular degeneration in CMT2D mice. Cell Death Dis. 11, 564. 10.1038/s41419-020-02798-y.

Sleigh, J.N., Tosolini, A.P., Gordon, D., Devoy, A., Fratta, P., Fisher, E.M.C., Talbot, K., Schiavo, G., 2020c. Mice carrying ALS mutant TDP-43, but not mutant FUS, display *in vivo* defects in axonal transport of signaling endosomes. Cell Rep. 30, 3655–3662.e2. 10.1016/j.celrep.2020.02.078.

Sleigh, J.N., Tosolini, A.P., Schiavo, G., 2020d. *In vivo* imaging of anterograde and retrograde axonal transport in rodent peripheral nerves. Methods Mol. Biol. 2143, 271–292. 10.1007/978-1-0716-0585-1_20.

Sleigh, J.N., Villarroel-Campos, D., Surana, S., Wickenden, T., Tong, Y., Simkin, R.L., Vargas, J.N.S., Rhymes, E.R., Tosolini, A.P., West, S.J., Zhang, Q., Yang, X.L., Schiavo, G., 2023. Boosting peripheral BDNF rescues impaired *in vivo* axonal transport in CMT2D mice. JCI Insight. 8, e157191. 10.1172/jci.insight.157191.

Sleigh, J.N., Weir, G.A., Schiavo, G., 2016. A simple, step-by-step dissection protocol for the rapid isolation of mouse dorsal root ganglia. BMC Res. Notes. 9, 82. 10.1186/s13104-016-1915-8.

Sleigh, J.N., West, S.J., Schiavo, G., 2020e. A video protocol for rapid dissection of mouse dorsal root ganglia from defined spinal levels. BMC Res. Notes. 13, 302. 10.1186/s13104-020-05147-6.

Snape, L., Sleigh, J.N., 2026. Junctions in Jeopardy: the neuromuscular junction is a selective pathological target in Charcot-Marie-Tooth disease. Mamm. Genome. 37, 72. 10.1007/s00335-026-10238-z.

Spaulding, E.L., Hines, T.J., Bais, P., Tadenev, A.L.D., Schneider, R., Jewett, D., Pattavina, B., Pratt, S.L., Morelli, K.H., Stum, M.G., Hill, D.P., Gobet, C., Pipis, M., Reilly, M.M., Jennings, M.J., Horvath, R., Bai, Y., Shy, M.E., Alvarez-Castelao, B., Schuman, E.M., Bogdanik, L.P., Storkebaum, E., Burgess, R.W., 2021. The integrated stress response contributes to tRNA synthetase-associated peripheral neuropathy. Science. 373, 1156–1161. 10.1126/science.abb3414.

Spaulding, E.L., Sleigh, J.N., Morelli, K.H., Pinter, M.J., Burgess, R.W., Seburn, K.L., 2016. Synaptic deficits at neuromuscular junctions in two mouse models of Charcot-Marie-Tooth type 2d. J. Neurosci. 36, 3254–3267. 10.1523/JNEUROSCI.1762-15.2016.

Tadenev, A.L.D., Hatton, C.L., Burgess, R.W., 2024. Lack of effect from genetic deletion of *Hdac6* in a humanized mouse model of CMT2D. J. Peripher. Nerv. Syst. 29, 213–220. 10.1111/jns.12623.

Theadom, A., Roxburgh, R., MacAulay, E., O’Grady, G., Burns, J., Parmar, P., Jones, K., Rodrigues, M., Impact CMT Research Group, 2019. Prevalence of Charcot-Marie-Tooth disease across the lifespan: a population-based epidemiological study. BMJ Open. 9, e029240. 10.1136/bmjopen-2019-029240.

Tinevez, J.Y., Perry, N., Schindelin, J., Hoopes, G.M., Reynolds, G.D., Laplantine, E., Bednarek, S.Y., Shorte, S.L., Eliceiri, K.W., 2017. TrackMate: An open and extensible platform for single-particle tracking. Methods. 115, 80–90. 10.1016/j.ymeth.2016.09.016.

Tosolini, A.P., Abatecola, F., Negro, S., Sleigh, J.N., Schiavo, G., 2024. The node of Ranvier influences the *in vivo* axonal transport of mitochondria and signaling endosomes. iScience. 27, 111158. 10.1016/j.isci.2024.111158.

Tosolini, A.P., Villarroel-Campos, D., Schiavo, G., Sleigh, J.N., 2021. Expanding the toolkit for *in vivo* imaging of axonal transport. J. Vis. Exp. 178, e63471. 10.3791/63471.

Trivedi, N., Jung, P., Brown, A., 2007. Neurofilaments switch between distinct mobile and stationary states during their transport along axons. J. Neurosci. 27, 507–516. 10.1523/JNEUROSCI.4227-06.2007.

Villarroel-Campos, D., Schiavo, G., Sleigh, J.N., 2022. Dissection, *in vivo* imaging and analysis of the mouse epitrochleoanconeus muscle. J. Anat. 241, 1108–1119. 10.1111/joa.13478.

Wei, N., Zhang, Q., Yang, X.L., 2019. Neurodegenerative Charcot-Marie-Tooth disease as a case study to decipher novel functions of aminoacyl-tRNA synthetases. J. Biol. Chem. 294, 5321–5339. 10.1074/jbc.REV118.002955.

Woschitz, V., Mei, I., Hedlund, E., Murray, L.M., 2022. Mouse models of SMA show divergent patterns of neuronal vulnerability and resilience. Skelet. Muscle. 12, 22. 10.1186/s13395-022-00305-9.

Ziermann, J.M., Boughner, J.C., Esteve-Altava, B., Diogo, R., 2021. Anatomical comparison across heads, fore- and hindlimbs in mammals using network models. J. Anat. 239, 12–31. 10.1111/joa.13409.

Zuko, A., Mallik, M., Thompson, R., Spaulding, E.L., Wienand, A.R., Been, M., Tadenev, A.L.D., van Bakel, N., Sijlmans, C., Santos, L.A., Bussmann, J., Catinozzi, M., Das, S., Kulshrestha, D., Burgess, R.W., Ignatova, Z., Storkebaum, E., 2021. tRNA overexpression rescues peripheral neuropathy caused by mutations in tRNA synthetase. Science. 373, 1161–1166. 10.1126/science.abb3356.

