## Supplementary for "The neuropathy-causing *GARS1*^Δ*ETAQ*^ mutation drives pathology in subsets of motor and sensory neurons in mice"

### Supplementary Figures

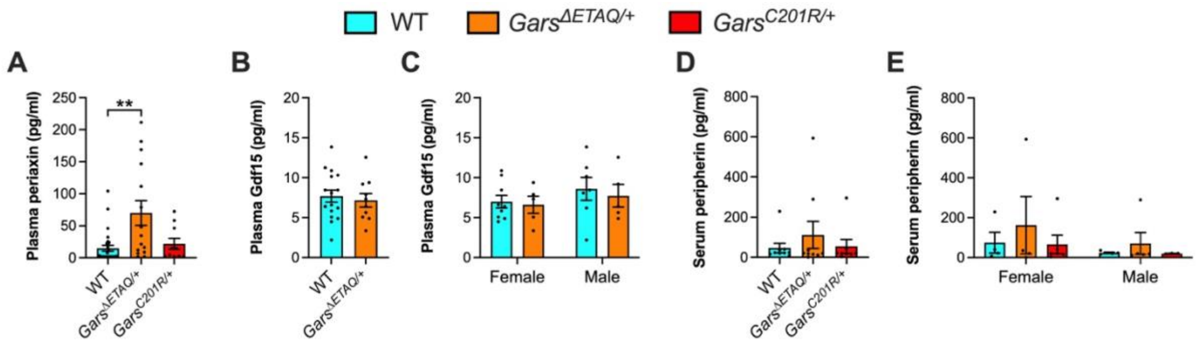


**Supplementary Figure 1. Gdf15 and peripherin levels are unaltered in *Gars*$^{\Delta ETAQ/+}$ blood at 3 months.** (**A**) Plasma periaxin levels are increased in *Gars*$^{\Delta ETAQ/+}$, but not *Gars*$^{C201R/+}$, mice relative to wild-type ($P = 0.008$ Kruskal-Wallis test). (**B-C**) Plasma Gdf15 levels are unaffected in *Gars*$^{\Delta ETAQ/+}$ mice (B, $P = 0.661$ unpaired *t*-test test; C, genotype $P = 0.606$, sex $P = 0.264$, interaction $P = 0.841$ two-way ANOVA). (**D-E**) Serum peripherin levels are also unaltered in both *Gars*$^{\Delta ETAQ/+}$ and *Gars*$^{C201R/+}$ mice (D, $P = 0.672$; E $P = 0.883$ Kruskal-Wallis test). $n$ = 10-29 (A), 10-16 (B), 5-9, (C), 8-9 (D) and 3-6 (E); $^{**}P < 0.01$ Dunn's multiple comparisons test.

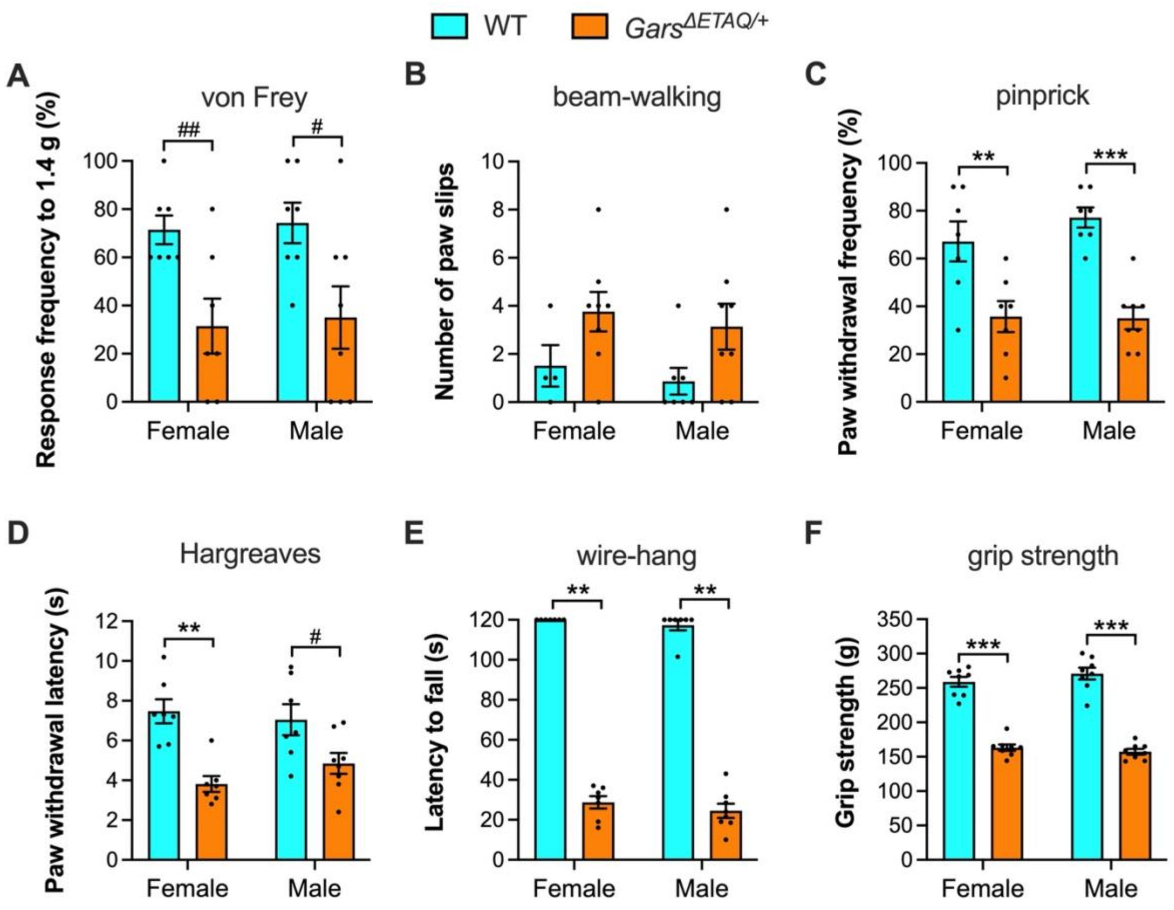


**Supplementary Figure 2. Sensory and motor behaviours are similarly impaired in female and male *Gars*$^{\Delta ETAQ/+}$ mice at 3 months.** (**A-F**) Female and male *Gars*$^{\Delta ETAQ/+}$ mice display equal deficits in mechanosensation (A, genotype $P < 0.001$, sex $P = 0.759$, interaction $P = 0.973$ two-way ANOVA), proprioception (B, genotype $P = 0.017$, sex $P = 0.477$, interaction $P = 0.992$ two-way ANOVA), mechanical nociception (C, genotype $P < 0.001$, sex $P = 0.452$, interaction $P = 0.386$ two-way ANOVA), thermal nociception (D, genotype $P < 0.001$, sex $P = 0.611$, interaction $P = 0.225$ two-way ANOVA), muscle endurance (E, $P < 0.001$ Kruskal-Wallis test), and maximal grip strength (F, genotype $P < 0.001$, sex $P = 0.644$, interaction $P = 0.171$ two-way ANOVA). $n$ = 7-8 (A, C-F) and 4-8 (B); $^{**}P < 0.01$, $^{***}P < 0.001$ Šídák's/Dunn's multiple comparisons test; $^{\#}$ $P < 0.05$, $^{\#\#}$ $P < 0.01$ unpaired *t*-test.

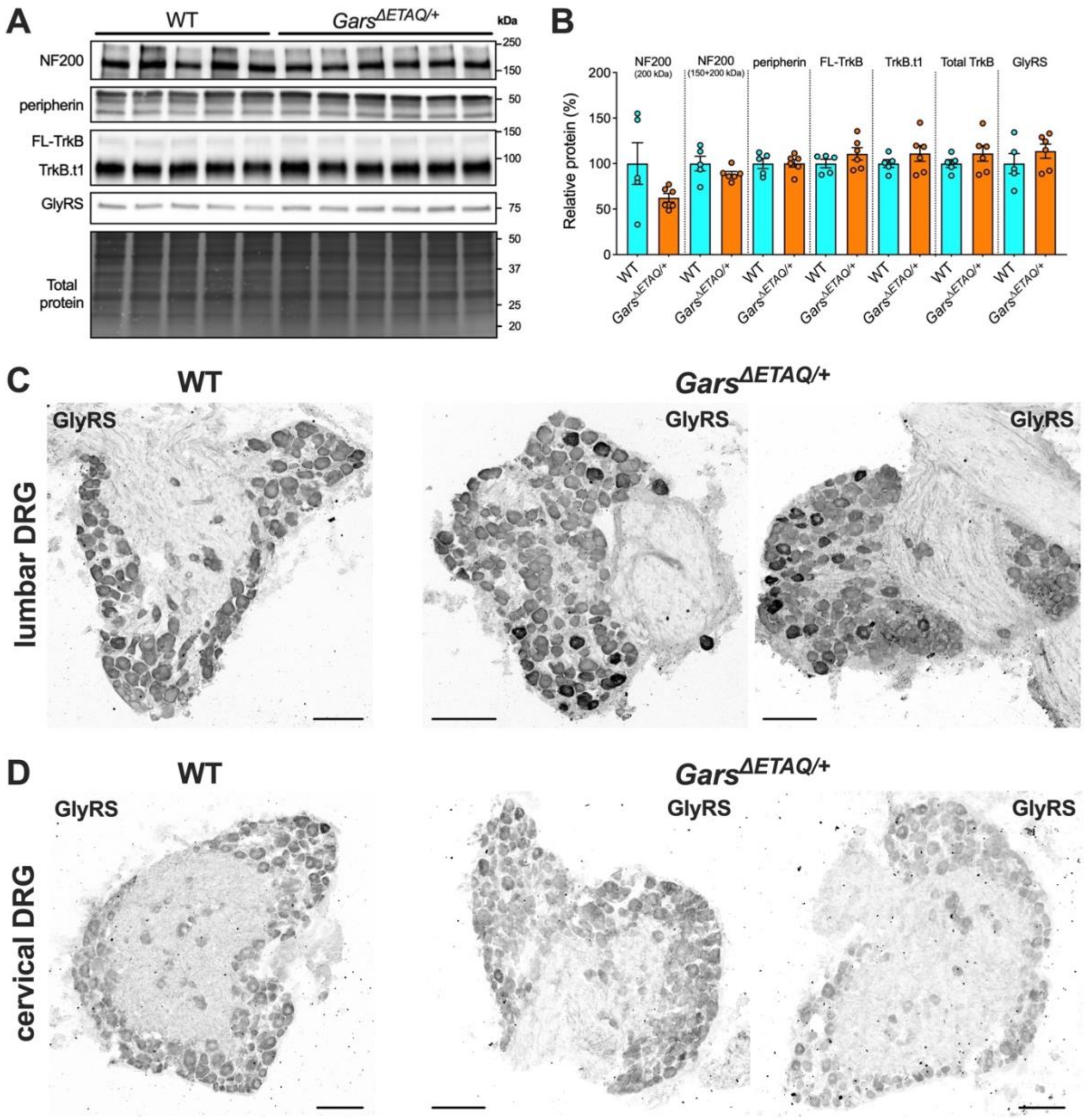


**Supplementary Figure 3. Sensory neuron subtypes are unaffected in *Gars*$^{\Delta ETAQ/+}$ cervical DRG.** (**A**) Western blots of C4-C8 DRG lysates from wild-type and *Gars*$^{\Delta ETAQ/+}$ mice aged P82-P143 probed with antibodies against several proteins that are altered in *Gars*$^{\Delta ETAQ/+}$ lumbar DRG (Sleigh et al., 2023). (**B**) Densitometric analyses show that, unlike in lumbar DRG, levels of NF200 (at 200 kDa, $P = 0.111$), total NF200 ($P = 0.178$), peripherin ($P = 0.969$), full-length TrkB (FL-TrkB, $P = 0.248$), truncated TrkB (TrkB.t1, $P = 0.313$), total TrkB ($P = 0.308$), and GlyRS ($P = 0.324$) remain unaltered in *Gars*$^{\Delta ETAQ/+}$ cervical DRG. (**C-D**) Representative single-plain confocal images of lumbar (C) and cervical (D) DRG sections from 3 month-old wild-type (left) and *Gars*$^{\Delta ETAQ/+}$ (right) mice stained for GlyRS. Scale bars = 100 μm. Genotypes were compared using unpaired *t*-tests; *n* = 5-6.

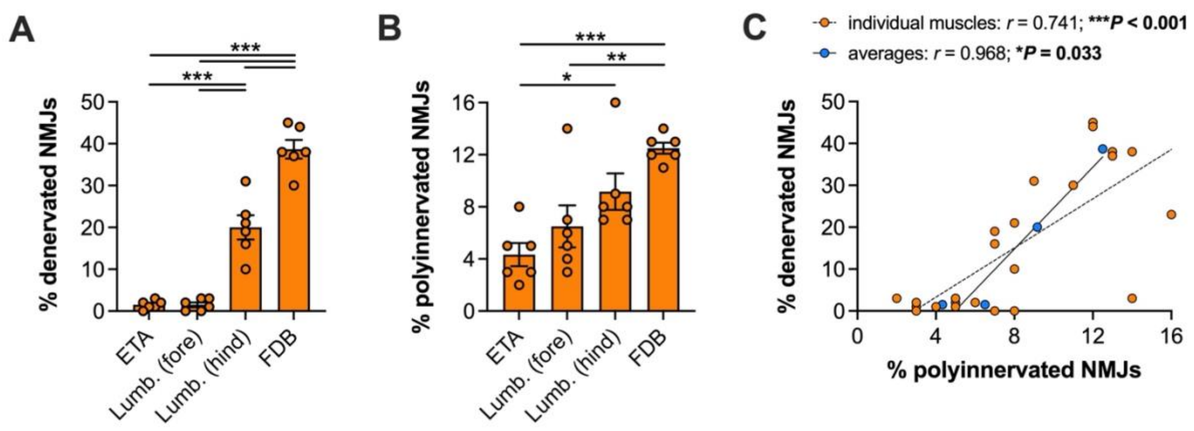


**Supplementary Figure 4.** ***Gars*$^{\Delta ETAQ/+}$ muscles display differential susceptibility to NMJ denervation and maturation impairment at 3 months.** (**A**) Hind-limb lumbrical and flexor digitorum brevis (FDB) muscles from *Gars*$^{\Delta ETAQ/+}$ mice display clear NMJ denervation (partial and vacant combined), whereas no such disruption in epitrocheloanconeus (ETA) or fore-paw lumbrical [*Lumb. (fore)*] muscles is observed ($P < 0.001$ one-way ANOVA). (**B**) NMJ maturation is differentially impacted across the assessed muscles, with the FDB and hind-paw lumbricals [*Lumb. (hind)*] displaying the greatest percentage of polyinnervated synapses ($P < 0.001$ one-way ANOVA). (**C**) NMJ denervation positively correlates with polyinnervation, both when individual muscles across mice (orange, $P < 0.001$) and average values for each muscle (blue, $P = 0.033$) are assessed. Data were analysed using Pearson's product moment correlation. *$P < 0.05$, **$P < 0.05$, ***$P < 0.001$ Tukey's multiple comparisons test; $n = 6$.

### Supplementary Tables

| Figure | Data | *n* | Sex (WT/*Gars*$^{\Delta ETAQ/+}$/*Gars*$^{C201R/+}$) | Age (d) |
|---|---|---|---|---|
| 1A-B | body weight | 12 | 12M12F/12M12F | 86-99 |
| 1C-D | plasma NfL | 11-14 / 5-8 | 6M8F/5M6F | 83-94 |
| 1E-F | plasma periaxin | 15-29 / 7-17 | 17M12F/7M8F | 71-101 |
| S1A | plasma periaxin | 10-29 | 17M12F/7M8F/3M7F | 71-101 |
| S1B-C | plasma Gdf15 | 10-16 / 5-9 | 7M9F/5M5F | 83-102 |
| S1D-E | plasma peripherin | 8-9 / 3-6 | 5M4F/5M4F/3M6F | 90-101 |
| 2A / S2A | von Frey | 14-15 / 7-8 | 7M7F/8M7F | 85-87 |
| 2B / S2B | beam-walking | 11-16 / 4-8 | 7M4F/8M8F | 83-85 |
| 2C / S2C | pinprick | 14-15 / 7-8 | 7M7F/8M7F | 85-87 |
| 2D / S2D | Hargreaves | 14-15 / 7-8 | 7M7F/8M7F | 84-86 |
| 2E / S2E | Wire-hang | 14-15 / 7-8 | 7M7F/8M7F | 86-94 |
| 2F / S2F | grip strenth | 8/16 | 8M8F/8M8F | 87-97 |
| 3B, E | lumbar DRG | 5 | 3M2F/3M2F | 89-93 |
| 3C | cervical DRG | 6 | 3M3F/4M2F | 89-93 |
| S3A-B | cervical DRG (WB) | 5-6 | 3M2F/3M3F | 82-143 |
| 4C | soleus muscle spindles | 5 | 2M3F/2M3F | 89-93 |
| 4D | biceps muscle spindles | 5 | 2M3F/4M1F | 98-101 |
| 5 | spinal cord motor neurons | 6 | 3M3F/4M2F | 89-93 |
| 6 / S4 | NMJs | 6 | 2M4F/3M3F | 92-98 |
| 7B-G | *in vivo* axonal transport (TA, GC) | 6 | 5M1F/3M3F | 89-95 |
| 7H-J | *in vivo* axonal transport (fore-paw) | 5 | 3M2F/0M5F | 89-95 |
| 7M | sciatic nerve WB | 7 | 3M4F/4M3F | 93-103 |
| 7N | sciatic nerve WB | 5-6 | 2M3F/3M3F | 93-103 |
| 7O-P | median/ulnar nerve WB | 8-9 | 4M4F/5M4F | 89-102 |

**Supplementary Table 1. Experimental sample sizes and mouse details.** *DRG*, dorsal root ganglion, *F*, female; *GC*, gastrocnemious; *M*, male; *n*, sample size; *NfL*, neurofilament light; *TA*, tibialis anterior; *WB*, western blot; *WT*, wild-type.

| Target | Sp. | Clonality | Company /Reference | Catalogue # | RRID | IF dilution | WB dilution |
|---|---|---|---|---|---|---|---|
| ChAT | Gt | Poly | Chemicon | AB144P | AB_2079751 | 1:250 | n/a |
| ERK1/2 | Rb | Poly | CST | 9102 | AB_330744 | n/a | 1:1,000 |
| p-ERK1/2 (T202/T204) | Rb | Poly | CST | 9101 | AB_331646 | n/a | 1:500 |
| GlyRS | Rb | Poly | Abcam | ab42905 | AB_732519 | 1:250 | 1:2,000 |
| Hook1 | Rb | Mono | Abcam | ab150397 | n/a | n/a | 1:1,000 |
| laminin | Rb | Poly | Sigma | L9393 | AB_477163 | 1:250 | n/a |
| neurofilament (2H3) | Ms | Mono | DSHB | 2H3 | AB_531793 | 1:250 | n/a |
| NF200 | Ms | Mono | Sigma | N0142 | AB_477257 | 1:500 | 1:1,000 |
| peripherin | Rb | Poly | Chemicon | AB1530 | AB_90725 | 1:500 | 1:1,000 |
| RILP | Rb | Poly | Abcam | ab140188 | n/a | n/a | 1:1,000 |
| Snapin | Rb | Poly | (Granata et al., 2008) | n/a | n/a | n/a | 1:1,000 |
| SV2 (pan) | Ms | Mono | DSHB | SV2 | AB_2315387 | 1:50 | n/a |
| TrkB | Rb | Poly | Merck | 07-225 | AB_310445 | n/a | 1:500 |

**Supplementary Table 2. Primary antibodies used in this study.** *ChAT*, choline acetyltransferase; *CST*, Cell Signaling Technology; *DSHB*, Developmental Studies Hybridoma Bank; *ERK*, extracellular signal-regulated kinase; *Gt*, goat; *IF*, immunofluorescence; *Ms*, mouse; *NF*, neurofilament; *Rb*, rabbit; *RILP*, Rab-interacting lysosomal protein; *RRID*, Research Resource Identifier; *Sp*, species; SV2, synaptic vesicle 2; *TrkB*, tropomyosin receptor kinase B; *WB*, western blotting.

| Target | Sp. | Conjugate | Company | Catalogue # | RRID | Dilution |
|---|---|---|---|---|---|---|
| Gt IgG | Dk | AlexaFluor555 | Life Technologies | A-21432 | AB_2535853 | 1:250-1,000 |
| Ms IgG | Dk | AlexaFluor488 | Life Technologies | A-21202 | AB_141607 | 1:250-1,000 |
| Ms IgG | Gt | AlexaFluor488 | Life Technologies | A-11001 | AB_2534069 | 1:250-1,000 |
| Ms IgG | Dk | AlexaFluor555 | Life Technologies | A-31570 | AB_2536180 | 1:250-1,000 |
| Ms IgG | Gt | AlexaFluor555 | Life Technologies | A-21424 | AB_141780 | 1:250-1,000 |
| Rb IgG | Dk | AlexaFluor488 | Life Technologies | A-21206 | AB_2535792 | 1:250-1,000 |
| Rb IgG | Dk | HRP | Bio-Rad | 1706515 | AB_11125142 | 1:3,000-5,000 |

**Supplementary Table 3. Secondary antibodies used in this study.** *Dk*, donkey; *Gt*, goat; *IgG*, immunoglobulin G; *HRP*, horseradish peroxidase; *Ms*, mouse; *Rb*, rabbit; *RRID*, Research Resource Identifier; *Sp*, species.

**Supplementary References**

Granata, A., Watson, R., Collinson, L. M., Schiavo, G., & Warner, T. T. (2008). The dystonia-associated protein torsinA modulates synaptic vesicle recycling. *J Biol Chem*, *283*(12), 7568–7579. https://doi.org/10.1074/jbc.M704097200

Sleigh, J. N., Villarroel-Campos, D., Surana, S., Wickenden, T., Tong, Y., Simkin, R. L., Vargas, J. N. S., Rhymes, E. R., Tosolini, A. P., West, S. J., Zhang, Q., Yang, X. L., & Schiavo, G. (2023). Boosting peripheral BDNF rescues impaired in vivo axonal transport in CMT2D mice. *JCI Insight*, *8*(9), e157191. https://doi.org/10.1172/jci.insight.157191
